# A Crossed Laser Phase Plate for CryoEM

**DOI:** 10.64898/2026.06.05.730245

**Authors:** Yue Yu, Anchi Cheng, Elizabeth Montabana, Noeli Paz Soldán, Eric S. Cooper, Jessie T. Zhang, Jeremy J. Axelrod, Petar N. Petrov, Amir Torkaman, Dylan Roof, Matthew Derstine, Bart Buijsse, Wim Hagen, Dari Kimanius, Shawn Zheng, Mykhailo Kopylov, Mohammadreza Paraan, Deepan Balakrishnan, Joshua Hutchings, Hang Cheng, Jonathan Remis, Ashwin Singh, Lothar Maisenbacher, Clinton S. Potter, Holger Müller, Bridget Carragher, David Agard, Pavel K. Olshin

## Abstract

The laser phase plate (LPP) enables phase-contrast imaging in cryogenic electron microscopy (cryoEM), enhancing image contrast without compromising high-resolution information. Here we report the implementation of a crossed laser phase plate (xLPP) comprising two optical cavities oriented orthogonally, installed in a ThermoFisher Scientific Krios G4 microscope equipped with a newly designed transfer lens module. We demonstrate the expected, strong contrast enhancement and stable, additive phase shifts of 90°, with a contrast transfer function (CTF) that closely matches theory. Single-particle analysis (SPA) of apoferritin, a standard benchmark sample, reached a resolution of 1.79 Å, demonstrating the system is capable of acquiring high-resolution cryoEM data. When imaging thick *E. coli* cells (∼350 nm), the xLPP enhances contrast and increases low-frequency template-matching signal. Together, these results establish the feasibility of the xLPP and highlight its potential for high-contrast, high-resolution cryoEM imaging of biological systems.

## Introduction

Transmission electron microscopy (TEM), especially cryogenic electron microscopy (cryoEM) and tomography (cryoET), is an essential tool for structural and cellular biology. ^1–4^ However, biological specimens are weak-phase objects that generate little intrinsic image contrast, limiting the achievable signal-to-noise ratio (SNR). Together with strict dose constraints, this restricts the resolution attainable for small or heterogeneous particles and complicates detection of molecules in crowded environments in tomograms. In principle, phase plates^5,6^ enable optimum SNR with weakphase objects^7^, but practical implementations in TEM, such as the Volta phase plate^8^, have thus far been limited by signal loss and instability. ^8–10^ Despite early promise, subsequent studies reported reduced final resolution in both single-particle analysis (SPA) of 167-kDa particles, and subtomogram averaging (STA) of ∼1-4 MDa particles, even with the increased contrast. ^11–13^

The continuous-wave laser phase plate (LPP)^14,15^ has been developed to address these limitations^16–19^ and has recently been shown to improve SPA resolution of small proteins such as hemoglobin (64 kDa)^20^. In this approach, the undiffracted portion of the electron beam is phase-shifted by 90° using a high-intensity laser field generated within a Fabry–Pérot cavity, reaching steady-state intensities of 350–400 GW/cm². ^16,19^ However, several aspects of the LPP could benefit from further hardware optimization. The high intra-cavity laser intensities required impose stringent demands on cavity construction and mirror performance, and also constrain the achievable focal radius of the intra-cavity laser beam ^21^, which in turn sets the cut-on spatial frequency of the phase plate. In addition, the LPP produces weak, unwanted “ghost” images, that can interfere with imaging, particularly in the presence of high-contrast objects, such as the specimen support film, or in heterogeneous samples. These limitations could, in principle, be alleviated by a crossed laser phase plate (xLPP), which uses two orthogonally-oriented laser beams ^22^. However, integrating and maintaining alignment of two intersecting cavities within the microscope column during data collection presents substantial technical challenges.

Here, we demonstrate a working prototype of a crossed laser phase plate (xLPP) that overcomes these challenges and enables stable operation during cryoEM data collection. Two orthogonal optical cavities are integrated as a compact unit within a customized Thermo Fisher Krios G4 microscope. The system produces additive phase shifts and imaging properties consistent with theoretical expectations. Using amorphous carbon films, we measure a stable phase shift of 89° ± 3°, achieved with ∼50% of the laser power per cavity compared to a single laser phase plate (sLPP), and observe the expected contrast transfer function modulation. We further validate the stability of the xLPP during biological data collection through single-particle analysis (SPA) of apoferritin and imaging of thick *E. coli* cells. We demonstrate the expected strongly enhanced low-frequency contrast and that automated repositioning of the electron beam can keep the electron beam aligned to both lasers throughout extended data collection sessions. Although apoferritin is not expected to benefit from phase-plate imaging^20^, it serves as a sensitive test for system performance. With the xLPP on, we achieve a resolution of 1.79 Å (FSC_0.143_), demonstrating the capability of the system for acquiring high-resolution SPA data. In a separate paired comparison using datasets acquired from the same grid (∼27,000 particles each), reconstructions reached 1.87 Å (xLPP on) and 1.93 Å (xLPP off). These results indicate that the optical cavities do not introduce aberrations or instability that compromise performance. Finally, Fourier ring correlation of 2D images from thick (∼350nm) *E. coli* samples, confirms enhanced low-frequency signal and demonstrates the feasibility of imaging in challenging, thick specimens. Together, these results establish the xLPP as a robust platform for cryo-EM and lay the groundwork for exploring its advantages across a range of applications in structural and cellular biology.

## Results

### Implementation of the crossed laser phase plate (xLPP)

The crossed laser phase plate (xLPP) consists of two independent, near-concentric Fabry-Perot optical cavities^16^ (Fig. 1a, b), where the power of the incoming laser beams (wavelength 1064 nm) is resonantly enhanced ∼8300 times. The two horizontally polarized laser standing waves have waist radii of ∼6 μm and ∼5 μm (1/e^2^ intensity), corresponding to numerical apertures (NAs) of 0.056 and 0.068, respectively (Extended Table 1). The lasers do not physically intersect but are displaced 100 μm along the electron beam propagation direction (Extended Data Fig. 1). This avoids possible laser interference while still being within the depth-of-field of the unscattered electron beam. Slight condenser astigmatism is used to tune the electron beam convergence along each laser axis to compensate for this offset. In the diffraction plane of the microscope, the two beam waists overlap within 20 µm, limited by mechanical tolerances. Nonetheless, the electron beam can be centered on this crossing point using electron beam deflectors.

**Fig. 1.**
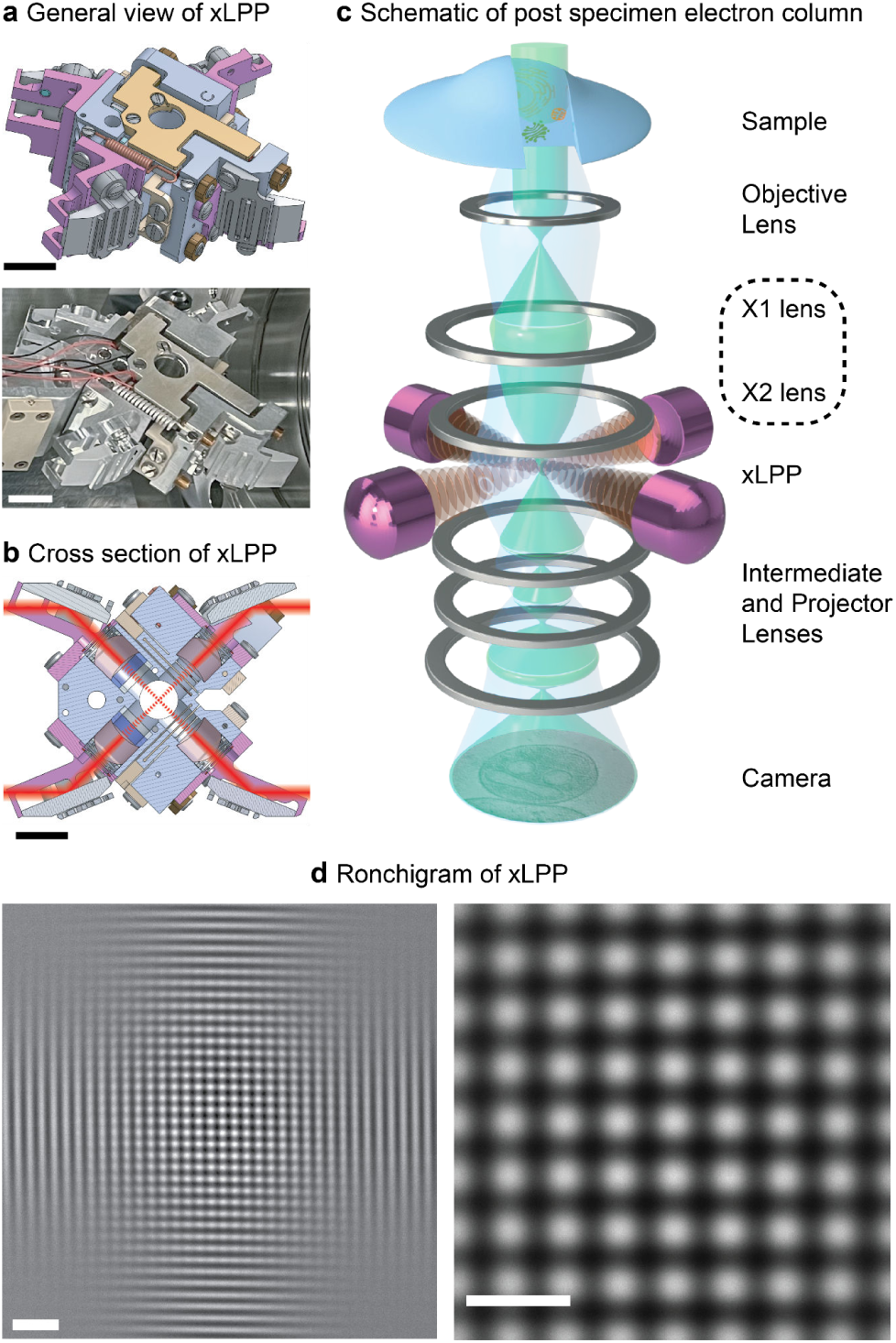
| Crossed laser phase plate (xLPP). **a**, CAD rendering (top) and a photograph (bottom) of a general view of the xLPP. Scale bars, 10 mm. **b,** Cross-section of the CAD model of the xLPP. Scale bar, 10 mm. **c,** Schematic of the post-specimen optical components in a microscope equipped with the xLPP. **d,** Far-field interference patterns of the e-beam diffracted by the laser standing waves, known as Ronchigrams, recorded at different magnifications. Scale bars, 2 µm (left) and 1 µm (right).

The xLPP is installed into a new model of TFS Krios G4 that accommodates a compartment for the phase plate (Fig. 1c). The microscope features additional “X-lens” (XL) modules (boxed in Fig. 1c, with a ray diagram in Extended Data Fig. 2) that transfers and magnifies the back focal plane to the xLPP plane, creates an image plane just upstream from the projection lenses, and enables aligning the e-beam to the laser beams. In XL mode, the nominal effective focal length (f), chromatic aberration coefficient (Cc), and spherical aberration coefficient (Cs) are *f* = 6.8 mm, Cc = 3.2 mm, and Cs = 2.7 mm, respectively. The shorter focal length and the carefully-designed X-lens unit improves aberrations compared to previous microscopes used with the LPP ^15,20^. The information limit measured by Young’s fringes was at 0.9 Å with xLPP installed (Extended Data Fig. 2). To characterize the laser fields introduced by the xLPP in the microscope, a Ronchigram^15^ can be recorded, which is formed when the electron beam is diffracted by the laser standing wave and interferes after focusing in the far field. Ronchigrams of the xLPP, recorded with the unscattered beam diverging at the phase plate plane, demonstrate close to orthogonal standing waves from both cavities of the xLPP (Fig. 1d).

### Phase shift and contrast transfer function (CTF) of the xLPP

To characterize the phase shift introduced by the xLPP, a series of amorphous carbon images was acquired with the e-beam positioned at different locations relative to the laser standing waves. Representative micrographs and measured phase shifts are shown where the unscattered e-beam is at an antinode–antinode intersection, an antinode–node saddle point, or a node–node intersection (Fig. 2a). When the e-beam is aligned to the antinodes of both standing waves, phase shifts of 86° ± 3° (n=6) were measured by GCtfFind^23^. The cavities were operating at circulating powers of 39 kW and 33 kW, with 3 hours pre-experiment idle time. Circulating power varied by only 0.2% over the duration of data collection as shown in Extended Data Fig. 3a.

**Fig. 2.**
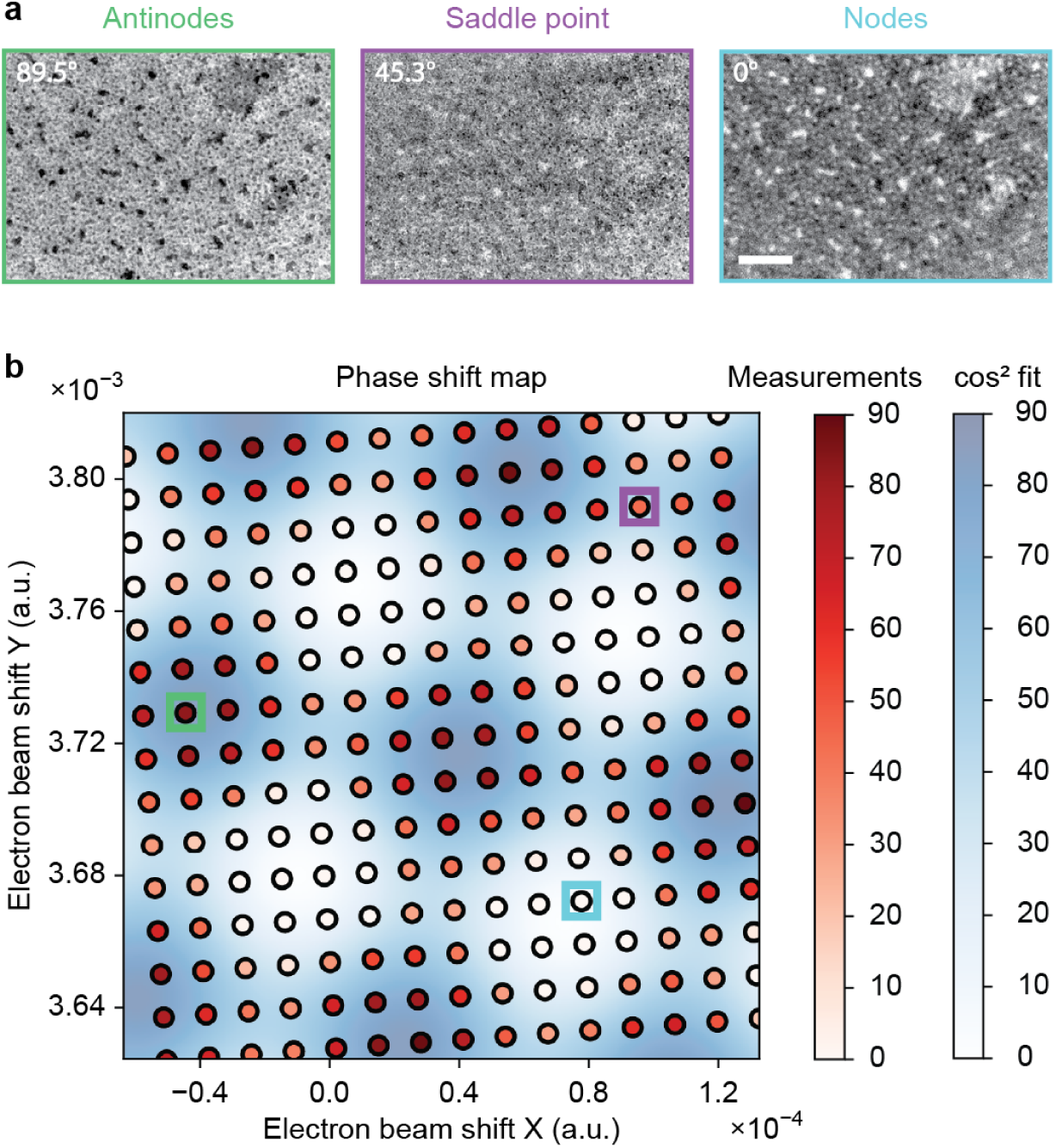
| Crossed laser phase plate (xLPP) phase-shift map. **a**, Representative images on amorphous carbon acquired with the electron beam centered at the intersections of the xLPP antinodes, the saddle point, and the intersection of nodes, illustrating contrast variations associated with different phase shifts as indicated in the micrographs. Scale bar, 50 nm. **b,** Two-dimensional phase-shift map showing experimental measurements (circles, white-red) overlaid on a fit (white-blue) consisting of two orthogonal squared-cosine (cos^2^) functions representing individual optical standing waves. The colored squares (green, purple, blue) correspond to the selected images shown in (**a**).

The measured phase shifts are plotted in Fig. 2b with circles (white-red) indicating the position of the e-beam at the phase plate plane as defined by the X-lens deflector settings. The data were fitted as the superposition of two squared-cosine (cos^2^) functions corresponding to the two laser phase plates (see Methods for the fitting details); the fitted map is shown as a continuous white-blue color map (Fig. 2b). Decomposition of the fitted map yields individual antinode phase shifts of 38° ± 2° and 46° ± 2°. The uncertainty arises from fitting the spatial phase shift map and does not account for CTF estimation uncertainty. In separate measurements when the unscattered e-beam is kept on the two xLPP antinodes, phase shifts measured with CTF on amorphous carbon were 88° ± 1° (n=7), and the expected phase shifts are 44° and 46°, estimated by each cavity’s circulating power and the NA (Ext. Data Fig. 4). Together these results demonstrate the phase shift introduced by the xLPP is an additive effect of the two laser phase plates, consistent with theory^22^.

**Fig. 3.**
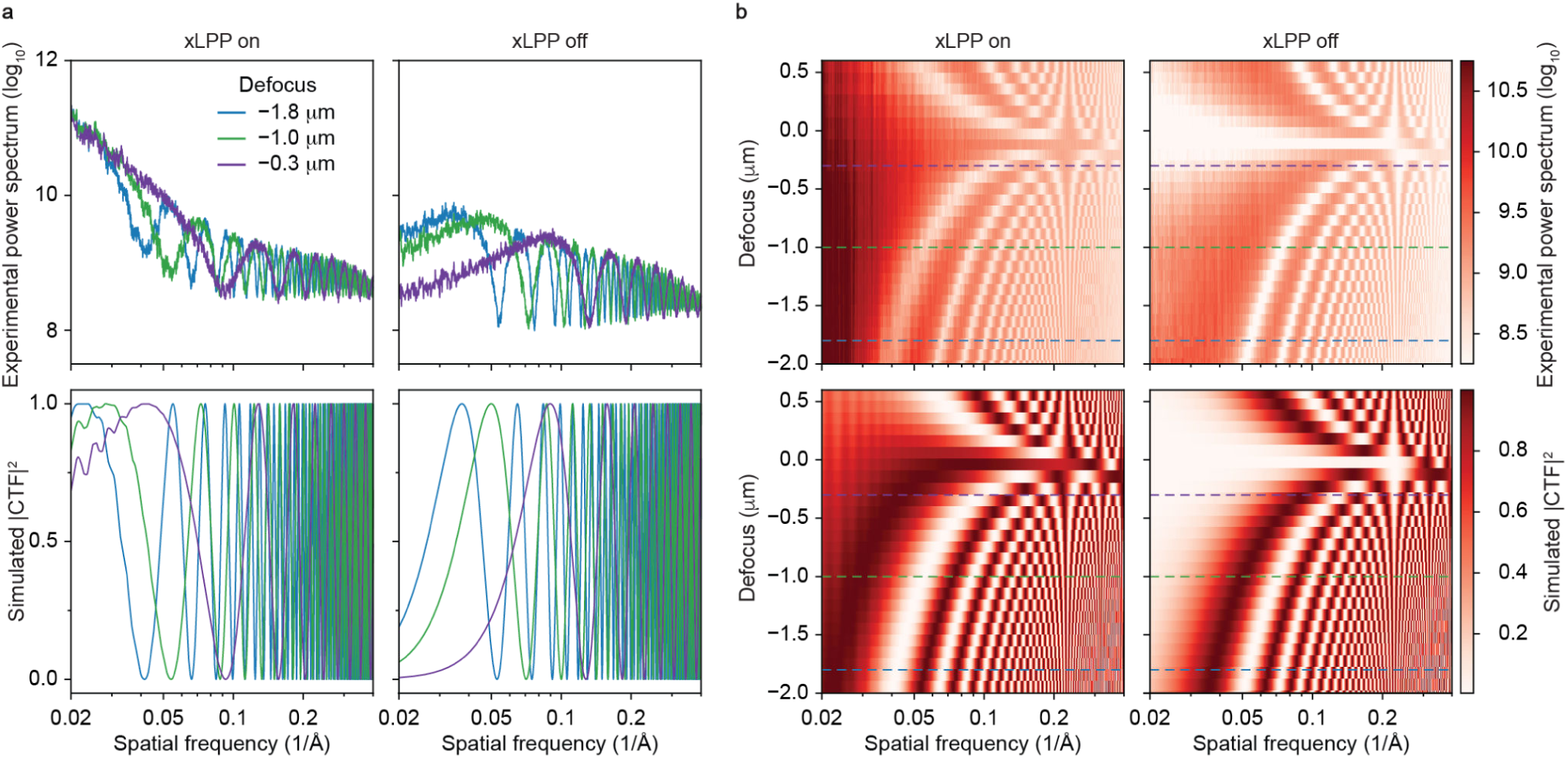
| Through-focus power spectra and CTF simulations with xLPP on and off. **a**, Radially averaged experimental power spectra (top) and simulated |CTF|^2^ (bottom) at three selected defocus values (−1.8, −1.0, and −0.3 µm; blue, green, and purple), with the unscattered beam aligned to the xLPP antinodes (left, phase shift 89° ± 3°) and with xLPP off (right). **b,** Through-focus radial averages of experimental power spectra (top) and simulated |CTF|^2^ (bottom) spanning the defocus range from –2 µm to 0.6 µm, for xLPP-on (left) and xLPP-off (right) conditions; colored dashed lines mark the three defocus values shown in **a**. The sample is amorphous carbon, and simulations use matching defocus and phase shift parameters.

**Fig. 4.**
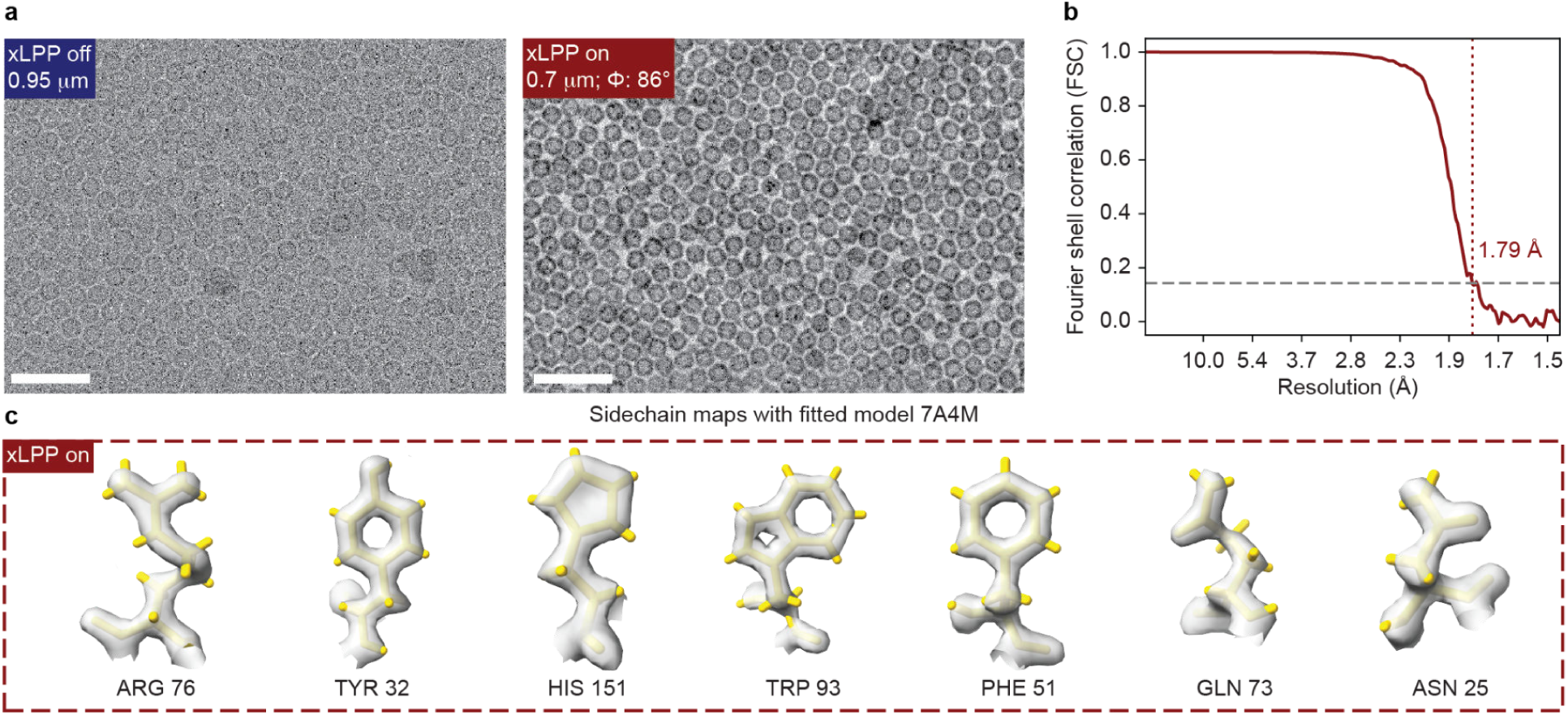
| Apoferritin single particle analysis with xLPP. **a**, Representative micrographs of apoferritin acquired without xLPP (0.95 μm defocus, left) and with xLPP (0.7 μm defocus, 86° phase shift, right), demonstrating a clear contrast enhancement due to the phase plate. The images are plotted using the same gray scale threshold percentile. Scale bar, 50 nm. **b,** Fourier shell correlation (FSC) indicates a resolution of 1.79 Å at the 0.143 threshold. **c,** Sidechain densities fitted with PDB 7A4M^1^ model confirm map quality at near-atomic resolution for xLPP on.

To verify that the xLPP modifies the contrast transfer functions (CTFs) as expected, a series of defocused images were recorded and compared to simulations with xLPP on and off. For the xLPP-on condition, the electron beam was aligned with the antinodes of the standing waves, yielding a measured phase shift of 89° ± 3°(n=21) across the defocus series. Because different sample regions were imaged in the xLPP-on and xLPP-off conditions, comparisons focus on the characteristic low-frequency features of the power spectra rather than absolute intensities. Fig. 3a shows radially averaged experimental power spectra and simulated |CTF|^2^ at selected defocus values, while Fig. 3b extends this comparison across a defocus range from −2 µm (underfocus) to +0.6 µm (overfocus) for xLPP-on (left) and xLPP-off (right) conditions. The sample was amorphous carbon, and the simulations use defocus and phase-shift parameters matched to experimental values. The experimental power spectra show the expected signal enhancement at low spatial frequencies and their agreement with CTF simulations confirms the predicted dependence on defocus and phase shift. Notably, detectable signal near zero defocus further demonstrates the feasibility of phase-plate imaging under near-focus conditions.

The (x)LPP introduces standing wave(s) at the back focal plane, such that scattered electrons passing through regions of laser intensity also acquire a phase shift. This effect is most pronounced at low spatial frequencies, where a larger fraction of scattered electrons traverse the laser intensity compared to higher frequencies. As a result, aligning the unscattered e-beam to the laser nodes also leads to additional contrast modification at low spatial frequencies (Extended Data Fig. 5). For the same reason, LPP CTFs present a standing wave pattern, and the impacted frequency region depends strongly on the effective focal length. With f = 6.8 mm, the standing wave pattern appears more magnified in the CTF than in previously reported LPP data^15^. With the xLPP configuration, the presence of two orthogonal lasers introduces a non-circular, fourfold-symmetric anisotropy in the power spectra, which is observed experimentally and reproduced by CTF simulations ^24^ (Extended Data Fig. 5). To evaluate the impact on CTF estimation, using fits performed with and without a cross-shaped mask that excludes the standing wave signal in the power spectrum, using GCtfFind^23^ (Extended Data Fig. 6). Excluding the standing wave pattern improved CTF fitting confidence and increased the estimated phase shift by ∼7°, while leaving the defocus estimate unchanged.

### Apoferritin single particle analysis (SPA) with and without xLPP

To demonstrate the feasibility of using the xLPP in the newly-designed microscope, and to ensure that neither introduces deleterious effects on imaging, we acquired several single-particle datasets of apoferritin. While the relatively large size and high symmetry of Apoferritin make it unlikely to benefit from LPP imaging²⁰, it provides an ideal test specimen for evaluating system performance, including electron optics and phase shift stability. During data collection, phase shift is affected by several factors that can change the position of the e-beam relative to the laser antinodes. These include changes in electron optics compounded by hysteresis, temporal (electron) beam-tilt drift ^1^, and thermal drift of the xLPP due to small environmental temperature fluctuations inside the microscope enclosure (Extended Data Figures 7-8). As a result, repositioning of the e-beam to both laser antinodes is required.^18,20^ An automated electron-laser alignment procedure for the xLPP was developed in Leginon ^25,26^, using a primary-reference Ronchigram image to correct the electron beam position using the X-lens deflectors (Extended Data Fig. 9). Phase shift stability across 1200 images from two datasets is summarized in Extended Table 2. In the current implementation, each exposure involves a stage shift, autofocusing, and beam alignment prior to image acquisition, resulting in an acquisition rate of ∼42 images per hour (∼1000 images per 24 hours).

Representative images with xLPP-off and xLPP-on are shown in Fig. 4a, using 0.95 µm defocus for xLPP-off and 0.7 µm defocus with 86° phase shift for xLPP-on. Despite the lower defocus, the xLPP-on image exhibits enhanced contrast (both images are displayed using the same grayscale threshold). To demonstrate high-contrast imaging at low-defocus, a dataset (Dataset A) was collected with xLPP-on with a mean defocus of 0.47 µm ± 0.18 µm, lower than typically used in conventional imaging. From 739 micrographs and 195,534 initial particle picks, 125,321 particles (64%) were retained, yielding a reconstruction at 1.79 Å resolution with a Guinier B-factor of 48 Å² (Fig. 4b). Representative side-chain densities with fitted models (Fig. 4c) confirm near-atomic resolution. Analysis of per-particle motion revealed drift up to ∼50 Å (Ext. Data Figure 10), likely arising from settling after movement between exposure targets and electron–laser alignment areas. This drift likely limited the overall resolution and contributed to an increased B-factor. To assess the impact, we split the dataset into four bins of equal particle sizes with increasing per-particle total drift. Both the Guinier B-factor and resolution show a clear correlation with drift, confirming its negative influence on data quality (Ext. Data Figure 10). To further verify that xLPP-on imaging does not compromise high-resolution information, a pair of datasets (Dataset B1 and Dataset B2) were collected on neighboring grid squares under identical optical alignment and similar imaging conditions. Processed using the same workflow, these datasets achieved comparable resolutions of 1.87 Å (xLPP-on, ∼0.4 µm mean defocus) and 1.93 Å (xLPP-off, ∼1 µm mean defocus) (Ext. Data Fig. 11). Acquisition conditions and dataset statistics for all three datasets are shown in Ext. Data Figure 12 and Extended Table 2. The comparable resolutions indicate that activating the phase plate does not introduce instability or aberrations that compromise high-resolution performance. Together these results demonstrate that the new microscope design with the xLPP installed is capable of high resolution SPA reconstruction.

### Enhanced contrast and low-frequency signal in thick *E. coli* cells with xLPP

A primary expected advantage of the phase plate is enhanced contrast when imaging complex *in situ* specimens. To illustrate this, the xLPP was used to image thick, un-milled *E. coli* cells (∼350 nm) overexpressing PP7 virus-like particles (VLPs) (Fig. 5a). Representative micrographs acquired at ∼1.6 µm defocus (xLPP off) and ∼1.1 µm defocus (xLPP on, 73° phase shift) are shown in Fig. 5a, with zoomed-in views in Fig. 5b; acquisition parameters are summarized in Table 1. “Ghost” images arising from Kapitza-Dirac diffraction of electrons by the laser standing wave^22^, are visible in features such as the membrane bilayer and ice contamination (black arrows in Fig. 5a). Additional examples are provided in Ext. Data Fig. 13. Across a range of defocus values and sample thicknesses, images acquired with the xLPP consistently exhibit improved visual contrast compared to those acquired without the phase plate.

**Fig. 5.**
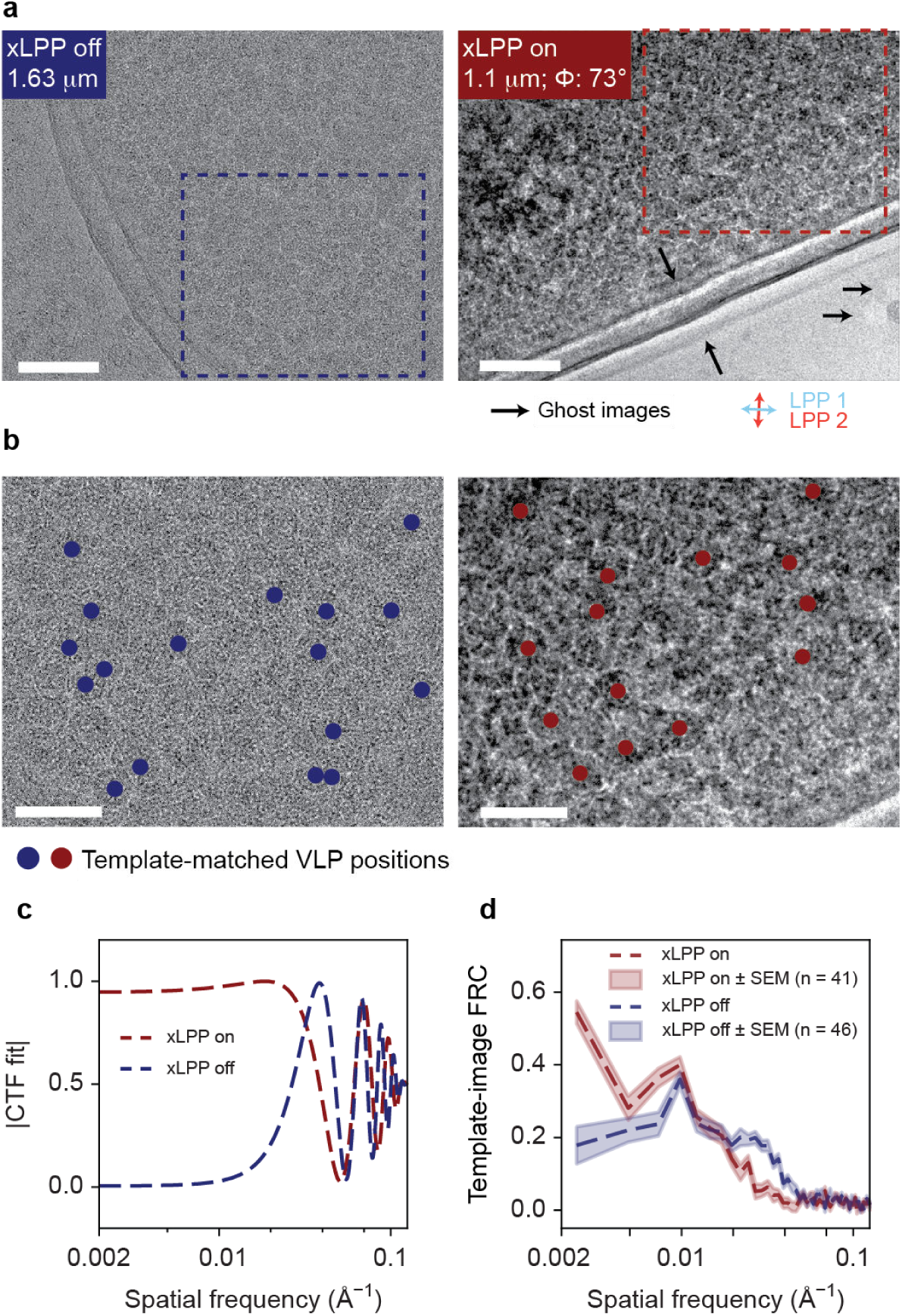
| xLPP improves contrast and template-matching of virus-like particles in thick *E. coli*. **a**, Representative micrographs of thick (∼350 nm) PP7-VLP-expressing *E. coli* acquired without (xLPP off) and with (xLPP on, 73° phase shift) the phase plate. Black arrows indicate ghost images of the membrane bilayer and an ice contamination. Sky-blue (LPP1) and bright red (LPP2) double-headed arrows indicate the propagation directions of each LPP. Scale bar, 100 nm. **b,** Zoomed-in views of the boxed regions in (a); blue and red circles mark representative template-matched VLP particle locations. Scale bar, 50 nm. **c,** CTF fits obtained with CTFFIND5 for both conditions (xLPP-on: 1.1 μm defocus, 73° phase shift; xLPP-off: 1.63 μm defocus). **d,** Mean Fourier ring correlation (FRC) between cropped particle images and the VLP template (xLPP-on, n=41; xLPP-off, n=46), with ±1 standard error of the mean (SEM) shaded, showing improved correlation at low spatial frequencies under the xLPP-on condition.

**Table 1.**
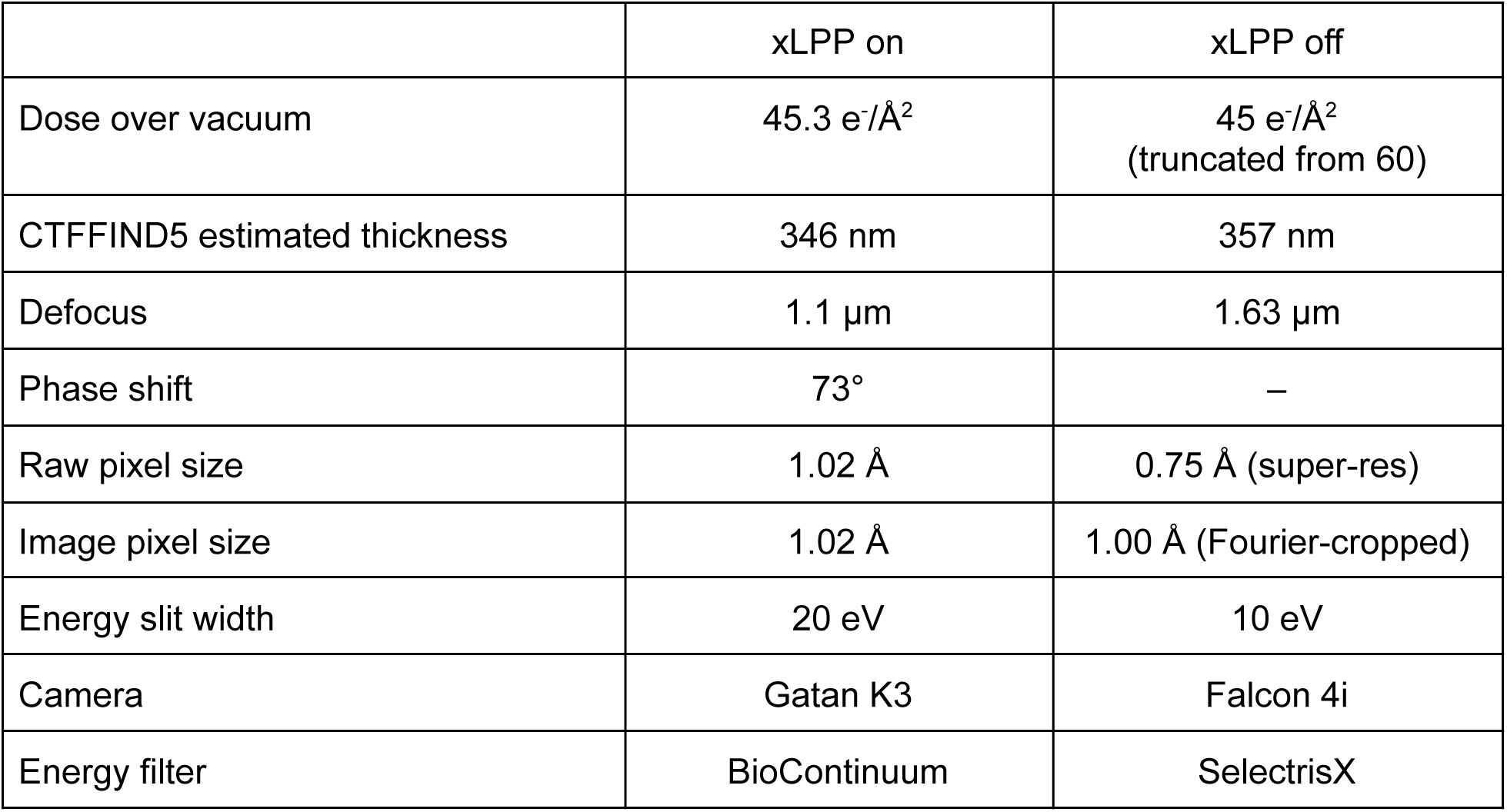
| Summary of the 2D *E. coli* imaging with and without xLPP.

Beyond visual contrast, template-matching (TM) signal was quantified using Fourier ring correlation (FRC) between particle images and the VLP templates at the orientations detected by TM. Cross-correlation thresholds were optimized separately for each micrograph and overall detectability was not directly compared. FRCs computed from 41 xLPP-on and 46 xLPP-off particles show enhanced signal in the xLPP-on condition at low spatial frequencies (<1/100 Å⁻¹), whereas the xLPP-off condition exhibits stronger signal in the intermediate frequency range (∼1/50–1/20 Å⁻¹) due to the choices of defocus values (Fig. 5d). These observations are consistent with theoretical expectations based on the corresponding CTFs (Fig. 5c): the xLPP enhances low-frequency signal below ∼1/30 Å⁻¹ (the crossover point), while xLPP-off images retain stronger signal between ∼1/30 and 1/18 Å⁻¹ (up to the first CTF zero). The FRC signal in the medium-frequency range may also reflect structural features of the PP7 VLPs, as different particle orientations exhibit distinct characteristics in this band (Extended Data Fig. 14). Nonetheless, the enhancement at low spatial frequencies can be robustly attributed to the phase plate.

To demonstrate the feasibility of xLPP tomography, we acquired a tilt series from the same apoferritin sample used for single particle analysis. The reconstructed tomogram shows the expected thin layer of apoferritin (< 25 nm ice thickness) without significant artifacts associated with suboptimal tilt series alignment (Extended Data Figure 18, Movie 1), demonstrating the feasibility of acquiring and aligning xLPP-on tilt series and tomogram reconstruction.

## Discussion

We demonstrate a crossed laser phase plate (xLPP) that delivers a stable phase shift up to 90°, implemented within a commercially-available xLPP-compatible ThermoFisher Scientific Krios microscope with minimal additional aberrations. The two cavities operate additively, each at approximately half the power required for a single LPP (sLPP), and produce contrast transfer function (CTF) modulations consistent with theoretical predictions. An automated alignment procedure maintains electron–laser overlap over extended data collection sessions (∼10–12 hours), supporting the practical usability of the system for routine cryo-EM imaging. The xLPP was applied to two representative specimen types: apoferritin by single particle analysis, and thick (∼350 nm) *E. coli* cells containing virus-like particles (VLPs). The apoferritin dataset reached 1.79 Å resolution, demonstrating high-contrast, high-resolution, at quite low defocus. In a paired comparison, reconstructions achieved 1.87 Å (xLPP-on) and 1.93 Å (xLPP-off), confirming that activation of the xLPP does not compromise high-resolution signal. Imaging of thick *E. coli* cells demonstrated a clear increase in contrast and enhanced low-frequency signal, quantified by Fourier ring correlation against VLP templates.

Apoferritin, owing to its large size and symmetry, is not expected to benefit substantially from phase-plate-induced contrast enhancement and is used here primarily as a benchmark for system performance. The observed B-factors are slightly higher than those reported for state-of-the-art systems ^1,27,28^, likely reflecting the current prototype data collection scheme in which the stage is repositioned after every exposure. This is not a fundamental limitation and will be improved in the future either by adopting optimized post-stage-shift settling times or implementing image-shift–based acquisition ^29^. For the *E. coli* data, the FRC analysis is based on a single paired laser-on/off comparison and should therefore be considered a proof of concept. A more comprehensive assessment will require larger datasets spanning a range of defocus values, sample thicknesses, and particle geometries. We observed that maintaining electron-laser alignment was more challenging for the *E. coli* samples than for apoferritin, beam deflection caused by charge accumulation in thicker specimens. In addition, we note that ghost images have been observed but quantitative characterization has yet to be undertaken.

The current implementation of the xLPP has room for further optimization. While the electron–laser alignment procedure was effective for the apoferritin dataset, it is not yet optimized to apply corrections only when necessary. A more robust strategy will require improved characterization and corrections of electron beam hysteresis, as well as enhanced beam-tilt stability and potentially reduced thermal drift of the laser cavities. Acquisition throughput, currently ∼42 images/hr, could be substantially increased by implementing beam-image shift data collection, which should be feasible with improved calibration of the X-lenses to minimize beam displacement in the back focal plane. Charge accumulation in biological specimens, which can act as an electrostatic lens and perturb electron–laser alignment, may be mitigated by conductive coatings^30^. For CTF estimation, excluding frequencies affected by LPP standing waves already improves fitting accuracy; in the future, full CTF deconvolution using an LPP-specific model may enable recovery of signal in these regions and potentially attenuate ghost image artifacts. Quantitative experimental validation of ghost attenuation relative to the sLPP, as well as direct measurement of the predicted reduction in cut-on frequency achieved by distributing power across two cavities, represent important near-term goals.

The results presented here open several avenues for broader applications. Recent studies using the sLPP have demonstrated improved performance in single-particle analysis of small proteins, including better motion correction, pose determination, and final resolution^20^. These advantages are expected to be even more pronounced in cryoET, where per-tilt contrast is inherently limited by dose fractionation. Systematic evaluation of xLPP performance in tomographic imaging, including contrast, tilt-series alignment, and subtomogram averaging, will be a natural next step as high-throughput cryoET workflows mature. Assessing the impact of cut-on frequency on tomographic imaging, and the relative performance of xLPP versus sLPP across different specimen types and imaging conditions, will require dedicated experimental and computational studies. As strategies to mitigate charging in lamellae continue to improve, extending phase-plate imaging to FIB-milled samples will further expand the range of accessible biological systems.

Ultimately, we believe the laser phase plate holds particular promise for structural biology for small proteins and for exploring in situ cellular contexts, where sample heterogeneity, low particle abundance, and crowded environments place the greatest demands on contrast and signal efficiency. After the planned future technical improvements outlined above, the laser phase plate promises to enable routine structural discovery in complex biological systems.

## Supporting information

Movie 1, high mag

Movie 1, low mag

## Methods

### xLPP design

The xLPP utilizes the same optical mirrors described previously^16^. Briefly, the radius of curvature of the concave is 10 mm and convex is 5 mm; nominal transmissivity of the high reflective (HR) coating: 0.01 %; the antireflective (AR) coating on the convex side of the mirror is 0.1% reflective.

The realization of the crossed cavities was done using the same basic geometry as the first LPP^16^. In order to fit two cavities in the same space as the single cavity, they were rotated 45° around the electron beam axis (Extended Data Figure 15). To direct the laser beams into the cavities and from the cavities to the output flange of the microscope, fold mirrors were added. The tip, tilt and separation adjustments were implemented using custom flexures to provide a travel range on the order of hundreds of micrometers and vibration and shock resistance. A comprehensive finite element analysis model (Solidworks Simulation Professional) of the flexures was developed to optimize their geometry and a high yield strength aluminum alloy (7075-T6) was used. The flexures were manufactured using electrostatic discharge machining, because it provides high accuracy in spatially restrained environments to achieve the required adjustment range of several hundred micrometers while maintaining acceptable stress levels.

The laser frequencies are stabilized to be resonant with the cavities using the Pound-Drever-Hall (PDH) technique. The optical systems to shape the beam and couple it to the cavity as well as to provide the PDH feedback and other diagnostic information closely followed the sLPP system ^16^, but was optimized to fit two sets of optical components within approximately the same volume. These systems were mounted on the vibration isolation system used by the microscope column.

During the xLPP-on operation, the scattered and absorbed light significantly heats up the system. To maintain the constant temperature of 35-36 °C during xLPP-on and xLPP-off, a resistive heating element has been installed on the arm that supports the xLPP at the center of the microscope. The voltage applied to the heating element is adjusted between different operational conditions.

### Characterization of optical cavities

The laser beam properties and relative position of the unscattered electrons relative to the laser position strongly affect the optical field intensity in the interaction region and therefore, the accumulated phase shift. Ideally, the electron beam should be centered at the antinode in both longitudinal and transverse directions relative to the propagation direction of the standing wave. The selected antinode should also be selected as close to the laser beam waist as possible to ensure the highest power density. In an ideal xLPP configuration, the two waists should overlap and this is determined by the location of the focal points of the fixed cavity mirrors. However, due to the manufacturing tolerances, this condition is not guaranteed. The xLPP was designed to have the waists of the two cavities within a Rayleigh range of the optical beams at the numerical aperture (NA) of 0.05, which is about 100 μm. To confirm the relative positions of the beam waists, a demagnified Ronchigram was reconstructed from a series of individual Ronchigrams to increase the field of view (Extended Data Fig. 1a). The images were first normalized by subtracting an image without the Ronchigram from images depicting laser standing waves, and then the absolute values were analyzed. This step ensured positive intensity between antinodes and nodes, which allowed using large integration windows to minimize signal to noise ratio in intensity profiles. Then, intensity profiles taken at different distances from the intersection position were fitted with Gaussian functions, and 1/e^2^ radii were extracted from the fits. The radii values *w*(*r*) were then plotted against the distance from the intersection *r* and fitted with a Gaussian beam distribution equation:

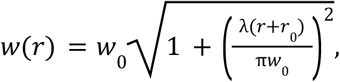

where 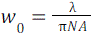 the beam waist (1/e^2^ radius), λ the laser wavelength (1064 nm), and *r*_0_ the beam waist offset from the intersection of the two beams. The fits (Extended Data Fig. 1b,c) demonstrate that the beam waists and distances to the overlap were around 6 μm and 1.3 μm for LPP1 and around 5 μm and 13 μm for LPP 2, respectively.

**Extended Table 1.** | Typical operation parameters for optical cavities.

| Parameter | LPP 1 | LPP 2 |
| --- | --- | --- |
| Length (mm) | 20 | 20 |
| Finesse | 13100 | 13100 |
| Effective numerical aperture | 0.056 - 0.055 | 0.068 - 0.064 |
| Waist ($\mu\text{m}$) | 6-6.1 | 5 - 5.3 |

The waists are determined by the radii of curvature (ROCs) of the cavity mirrors, the laser wavelength, and the NA of the cavity, which depends from the distance to concentricity. During normal operation conditions, the NA is the only varying parameter, and typically it is kept above 0.05, which is achieved at a distance to concentricity below 3.4 μm for the mirrors with the ROC of 10 mm and the wavelength of 1064 nm. The optical cavity, formed by mirrors that deviate slightly from a perfect spherical shape, exhibits a small amount of astigmatism. In a near-concentric cavity, however, even this small astigmatism can produce substantial differences in the beam waist along different axes of the laser beam. This difference can exceed 1 μm, corresponding to about 20% for a beam waist of roughly 5 μm. Therefore, determining the orientation of the long and short axes relative to the electron beam is important for a more accurate estimate of the phase shift.

Due to the large difference in the NAs caused by the astigmatism, the first order cavity modes TEM_01_ and TEM_10_, which are aligned along two principal axes, can be well separated spectrally (Extended Data Fig. 16a). This makes it possible to lock the laser frequency to these cavity modes and characterize them independently. Extended Data Fig. 16b,c shows Ronchigrams of the TEM_01_ and TEM_10_ modes, but only one of them exhibits a zero intensity region (Extended Data Fig. 16c). This indicates that the line connecting the centers of the bright parts of the first-order mode with the higher NA (marked in green) is oriented nearly parallel to the electron beam, whereas the corresponding line for the other mode (marked in red) is nearly orthogonal to it. The intensity profiles across the laser beam match the intensity profiles of camera images of the corresponding modes, which indicates that the camera images may be used for the preliminary estimation of the astigmatism orientation before the installation of the xLPP inside a TEM.

The expected phase shift stability was estimated using the circulating power of the beam and the beam radii at the beam intersection (Extended Data Fig. 4). The circulating power was calculated using a calibrated power meter that measures the power of the laser beam transmitted through the optical cavity and the total losses of the cavity mirrors measured using the cavity ring down (CRD) method. The beam radii at the intersection were estimated using the method similar to that described in Extended Data Fig. 1. A series of Ronchigrams was recorded overnight with a ∼6 minute interval between the images. After that, the waists for both standing waves as well as the beam radii at the intersection position were determined at each time. To calculate the linear power density, the circulating power was divided by the beam radius. As the circulating power is recorded at ∼10 Hz rate, the beam waist values were linearly interpolated for time points for which no corresponding image was recorded. Sudden changes in the circulating power observed around 18:00 and 03:00 are associated with cavity unlocking events caused by filling the microscope systems with liquid nitrogen (Extended Data Fig. 4c). After the process finished, the laser frequency was successfully locked to that of the optical cavity for both resonators. The estimated phase shift histograms demonstrate that the phase shift stays extremely stable (within ±0.3° as determined as a standard deviation of Gaussian fits of the histograms) for an extended period of time. The standard deviation for the mean power density appears significantly larger than that of the expected phase shift as it is significantly affected by the “unlocking” event, during which the circulating power was around zero (Extended Data Fig. 4b). For the expected phase shift, the standard deviation was calculated from the Gaussian fit. We note that the phase shift measured from a power spectrum of a micrograph of amorphous carbon recorded at these parameters, demonstrates a phase shift of 88° ± 1° as determined using the method described in Extended Data Fig. 6, which is in good agreement with the expected phase shift based on the sum of linear power densities (90.6° ± 0.3°).

### xLPP induced phase shift as a function of e-beam position at the phase plate plane

To characterize the phase shifts generated by the xLPP, the unscattered electron beam was purposely deflected across a 2D grid at the phase plate plane using the X-lens deflectors. 225 images of amorphous carbon were recorded across a 15 × 15 raster grid, and the phase shift of each image was estimated by GCtfFind^23^, with measurements close to 180° converted to 0°, (Fig. 2b, white-red circles). The phase shift map was modelled as the superposition of two squared-cosine functions, one per LPP:

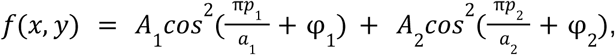

where *p*_1_ = (*x* − *r*_x_)*cos*θ + (*y* − *r*_*y*_)*sin*θ and *p*_2_ = (*x* − *r*_x_)*cos*(θ + δ) + (*y* − *r*_*y*_)*sin*(θ + δ) are projected coordinates along the two lattice directions, (*r*_x_, *r*_*y*_) is a position reference offset, θ is one fringe’s orientation and δ is the angle between the two near-orthogonal fringes. *A*_1_ and *A*_2_ are amplitudes, φ_1_ and φ_2_ are phase shift offsets, and *a*_1_ and *a*_2_ are periodicities. The model was fitted to the scattered measurements by nonlinear least squares (L-BFGS-B), using a U-shaped weighting that emphasizes peak and trough values, with a 3× penalty for under-predictions to discourage underestimation of peak phase shifts. The fitted map is shown in Fig. 2b as the continuous white-blue colourmap; decomposition of the fit yields individual contributions of 38° ± 2° (*A*_1_) and 46° ± 2° (*A*_2_) to the peak antinode phase shift; reported uncertainties are 1σ, derived from the fit covariance matrix

### Experimental CTF validation against theory

Through-focus image series of an amorphous carbon film were acquired under xLPP-on and xLPP-off conditions, on different regions of the grid for each case. For both xLPP-on (unscattered beam aligned to the laser antinodes) and xLPP-off conditions, images were collected from a nominal defocus of approximately −2.0 µm (underfocus) to +0.6 µm (overfocus) using the same nominal defocus step size between consecutive exposures. The defocus value of every underfocused image was estimated by GCtffind^23^, and the defocus values for on-focus and overfocus images were obtained by linear extrapolation from the calibrated underfocus values. For xLPP-on, phase shifts measured from the underfocus images were 89° ± 3°. Radially averaged power spectra were computed as the squared modulus of the 2D FFT, averaged into 4000 radial bins, with the *log*_10_ transform applied for visualisation. Radial profiles were stacked along the defocus axis, sorted from most underfocused to most overfocused, and clipped to the spatial-frequency range 0.02–0.40 Å⁻¹ for Fig. 3.

CTF simulations were generated for the same defocus range and a similar peak phase shift as the experimental data. For xLPP-on, the phase shift introduced by the xLPP standing wave at the back focal plane was computed from an analytical model of two orthogonal Gaussian standing-wave cavities ^22^. For simplicity, both cavities were modeled with identical power, numerical aperture (NA = 0.05), and wavelength (1064 nm), yielding an additive peak phase shift of π/2 (90°). The xLPP-on case was modeled with the electron beam aligned to the antinodes of both laser standing waves. The 2D phase shift was azimuthally averaged about the unscattered-beam position to produce a 1D radial profile φ(*s*) representing the orientation-averaged phase shift seen by diffracted electrons at spatial frequency *s*. The 1D CTF was computed with φ(*s*) as an additional phase shift, and |CTF|^2^ was plotted in Fig. 3. Since the anisotropic modulations caused by the xLPP phase shift are averaged out in the 1D radial profiles, a two dimensional comparison is required.

When the electron beam passes precisely through an antinode or node of the standing wave, the xLPP phase produces symmetric modulations in the CTF; any slight misalignment causes asymmetric modulations. However, since the detector records intensity, the resulting image is real-valued and its Fourier transform obeys Hermitian symmetry. As a result, Hermitian symmetry folds the asymmetric components in the power spectrum, rendering them indistinguishable from the symmetric components. This makes a direct comparison between the full complex CTF applied during the image formation and the experimental power spectrum insufficient for validation.

Hence, we compare the 2D power spectra of images simulated with a forward model against those of experimental images, where both have undergone the same intensity detection process (Extended Data Fig. 5). The xLPP phase shifts are simulated using the torch-CTF package^24^, with laser wavelength (1064 nm), NA (0.05), and an effective focal length (6.8 mm). The simulated images are generated using a single-slice weak-phase numerical phantom as a specimen modulated by a complex CTF that includes defocus, aberrations and the xLPP phase shift. The simulated and experimental log power spectra show close agreement in the non-circular, four-fold symmetric structure and zero crossings arising from the crossed geometry of the xLPP (Extended Data Fig. 5). Since the simulated phantom does not replicate the exact specimen and we do not account for beam incoherence effects, deviations in the spread of the central bright region are expected.

### Drift of electron-laser relative position over a 31-hour acquisition

Electron-laser relative position is the primary cause of instability of LPP’s phase shift. Hysteresis from changes in electron optics (e.g. beam diameter, magnification, or beam blanking) can cause the e-beam’s position at the phase plate plane to vary. The e-beam also undergoes a temporal beam-tilt drift, as previously reported on similar microscopes ^1^. Additionally, environmental temperature changes can cause mechanical motion of the laser phase plate.

Extended Data Figure 7 and Extended Data Figure 8 isolate the effect of environmental drift by continually acquiring Ronchigrams over a cross-grating grid over 31 hours without changing stage position, beam diameter or magnification. The images capture the movement of the laser through a far-field interference pattern of the xLPP fringe (blue box). It also measures the movement of the electron beam footprint by choosing magnification and beam diameter where the electron beam edge (orange circle) was in the field of view (FOV), and measures the movement of sample / stage with imaging in-focus a cross-grating grid with a latex bead (green box) in FOV. Images are taken every 5 min excluding the time during liquid nitrogen refill and/or cold field emission gun flash.

Images taken at time points are analyzed as follows. Edge detections are applied to the beam foot print and these part circles were fitted to obtain the center of the circle. The latex bead is boxed out and binary-threshold applied to highlight the bead and center of mass calculated. The crossed region of the LPP Ronchigram was boxed out to exclude the bead and edge of the electron beam. It was then analyzed using cross-correlation with other time points with a correction for global correlation profile.

We observe that nitrogen dewar refills are the primary source of environmental perturbation, leading to temperature fluctuations of 0.2 °C. The measured position of the laser tracks temperature fluctuations with a delay of ∼1 hr. We measured faster drift (up to ∼15 nm/min) during and 1-2 hours after liquid nitrogen (LN2) refill, and otherwise slower drift (2.1 nm/min ± 1.3 nm/min) for the laser standing waves (Extended Data Fig. 8) in the xLPP plane. For reference, the node-to-node distance for the laser standing waves is 532 nm. The observed drift is consistent in direction with the expected differential expansion between the aluminum support rod and steel vacuum chamber. Residual drift that is not explained by the temperature cycles is approximately 100 nm over 10 hours and could be explained by a beam tilt of just 15 microradians. This drift is well within the typical values seen during high-resolution cryo-EM data collection, with some studies observing beam tilt, as measured by coma, drift at a rate of tens of microradians per hour ^1^.

### Leginon-based auto electron-laser alignment procedure

Due to a combination of electron beam hysteresis, electron beam tilt drift, and the laser beam temperature drifts, alignment of the electron beam to the laser beam is performed during data collection. The electron-laser automatic alignment is based on Ronchigrams, as illustrated in Ext. Data Fig. 9. At the beginning of data collection, the user adjusts the electron beam in the transferred back focal plane to optimize the phase shift under data acquisition conditions. A reference image is acquired to record the Ronchigram at optimal alignment. During data collection, alignment-check Ronchigrams are periodically acquired from the alignment area and compared to the primary reference to maintain optimal electron–laser alignment. The reference can be acquired either on supporting foil or over the hole, on the sample. Dataset A (Fig. 4) was acquired with reference over the hole, on the sample, Dataset B1 was acquired with reference on carbon supporting foil.

### Sample preparation for apoferritin and *E. coli* overexpressing PP7 VLPs

For the apoferritin sample used in figure 4 and Ext. Data Fig. 11, mouse Heavy Chain Apoferritin^27^ was purified in house and frozen down for storage in –80. Subsequently aliquots (10+ mg/ml) were thawed on ice and 1 mM DTT was added immediately before plunge freezing 3 ul on either Ultraufoil 300 1.2/13 (Dataset A), Quantifoil Au 200 1.2/1.3 (Datasets B1 and B2) grids on a ThermoFisher Vitrobot MKIV. Temperature was set to 10C, 100% Humidity, 30s wait time, Blot Force 0, varying blot times.

The PP7 virus-like particles (VLPs) were produced by transforming BL21(DE3) *E. coli* with PP7-dimer plasmids and growing the culture to an optical density (OD₆₀₀) of 0.6 before inducing protein expression with IPTG. Cell pellets were collected by centrifugation and stored. For cryoelectron microscopy sample preparation, a pellet was resuspended in LB broth and adjusted to an OD₆₀₀ of 0.6, a concentration found to be optimal for forming a thin monolayer suitable for imaging without the need for focused-ion beam milling. Quantifoil R 2/1 Au 200 mesh grids were glow-discharged in a Ted Pella Pelco easiGlow at 15 mA for 45 seconds. 4 μL of the sample was applied to the grid, blotted for 4 seconds, and plunge-frozen in liquid ethane using a Leica GP2 plunge-freezer under controlled conditions (90% humidity, 4°C chamber temperature).

### Apoferritin SPA with and without XLPP

For all three datasets (Datasets A, B1, and B2), stage movement was used for navigation between targets. The cavity circulating power during Dataset B1 acquisition is shown in Ext. Data Fig. 3b; the sharp drops were unlocking events caused by stage movements and LN2 refill, but quickly automatically relocked after the stage stops. Extended Table 2 summarizes the collection conditions.

Micrographs were curated based on cryoSPARC metadata. For the xLPP-on datasets (A and B1), micrographs with defocus values below 1 µm and CTF fit confidence better than 4.5 Å were retained, and an upper bound of 100° was applied to phase shift values based on cavity power and numerical aperture (NA). For the xLPP-off dataset (B2), micrographs were selected with CTF-estimated defocus values between 0.6 and 1.5 µm and CTF fit confidence better than 4.5 Å. For all three datasets, outliers in relative ice thickness, defocus tilt and range, and micrograph intensity were removed. The distributions of phase shift, defocus, and CTF resolution after curation are shown in Extended Table 2 and Ext. Data Fig. 10. For defocus, CTF fit resolution and phase shift In Extended Table 2, mean ± standard deviation is reported. Additionally, as an option in cryoSPARC, phase shift values measured with CTFFIND4 are also reported.

For all three datasets, particles were picked using a blob picker, and good particles were selected by 2D classification based on visible secondary structure. An initial model was built from 10,000 particles and subsequently refined using all selected particles. After refinement, particles were further curated based on per-particle scale in cryoSPARC. The final refinement included optimization of per-particle defocus and per-group CTFs, along with correction for spherical aberration, tetrafoil, anisotropic magnification, and Ewald sphere (EWS) effects. A final step of reference-based motion correction was then applied using the refined map as reference, estimating per-particle trajectories from the refined poses and calculating empirical dose weights from the Fourier cylinder correlation (FCC) to optimally account for radiation damage. The Rosenthal–Henderson (RH) B factor was calculated from a ResLog analysis with symmetry and EWS correction, starting from 100 particles. For Dataset B2, the final particle count was randomly subsampled after per-particle scale curation to match the number of particles in Dataset B1 for a direct comparison.

Dataset B1 (xLPP-on) had a substantially lower final particle retention rate (∼25%) than the other two datasets, likely due to the use of two exposures per hole: stage-shift targeting inaccuracies may have increased the fraction of mis-targeted exposures landing on the carbon support, and targets placed near the supporting foil are more prone to thicker ice and overlapping particles. Dataset A showed worse B factors than the paired Datasets B1 and B2, which may reflect the shorter post-movement wait time used during its collection. The wait time was the same for Datasets B1 and B2, collected from the same grid, serving as the xLPP-on versus xLPP-off comparison.

**Extended Table 2.** | Summary of the apoferritin SPA with and without xLPP. * The pre-exposure wait is the dwell time after the stage moves to the exposure area following autofocusing and auto-Ronchigram alignment; it excludes the time required for those alignment steps themselves.

|  | Dataset A<br>(xLPP-on) | Dataset B1<br>(xLPP-on) | Dataset B2<br>(xLPP-off) |
| --- | --- | --- | --- |
| Grid | Ultrafoil 300<br>1.2/13 | Quantifoil Au 200<br>1.2/1.3 | Quantifoil Au 200<br>1.2/1.3 |
| Dose | 42 e<sup>-</sup>/Å<sup>2</sup> | 49 e<sup>-</sup>/Å<sup>2</sup> | 43 e<sup>-</sup>/Å<sup>2</sup> |
| Number of frames | 34 | 34 | 34 |
| Pre-exposure wait time* | 2.5-5 s | 7.5 s | 7.5 s |
| Number of exposure per hole | 1 | 2 | 1 |
| Micrographs | 739 | 476 | 225 |
| Micrographs curation | 599 (81%) | 350 (74%) | 177 (79%) |
| Defocus | (0.47 ± 0.18) μm | (0.43 ± 0.17) μm | (0.98 ± 0.19) μm |
| CTF fit resolution | (2.75 ± 0.35) Å | (2.79 ± 0.45) Å | (2.91 ± 0.29) Å |
| Phase shift | (53 ± 13)° | (69 ± 14)° | - |
| Phase shift (CTFFIND4) | (68 ± 20)° | (84 ± 13)° | - |
| Initial particle picks | 195,534 | 112,610 | 76,026 |
| Post 2D classification picks | 141,855 (73%) | 37,942 (34%) | 57,635 (76%) |
| Per-particle scale curation | 126,270 (64%) | 27,730 (25 %) | 38,017 (50%) |
| FSC map resolution (Å) | 1.79 Å<br>(125,321 particles) | 1.87 Å<br>(27,204 particles) | 1.93 Å<br>(27,202 particles) |
| Rosenthal–Henderson B-factor (Å<sup>2</sup>) | 86 | 61 | 78 |
| Guinier B-factor (Å<sup>2</sup>) | 48 | 40 | 45 |
| EMDB | EMD-77042 | EMD-77043 | EMD-77047 |

### 2D template matching signals in *E. coli* cells

For xLPP-off, images were collected in super-resolution mode at 0.75 Å/pixel and Fourier-cropped to 1.00 Å/pixel; for xLPP-on, the image pixel size was 1.02 Å. For xLPP-off, the total dose over all frames was 60 e^-^/Å^2^ and 45 frames were used to match the total dose of xLPP-on, which is 45.3 e^-^/Å^2^. All subsequent operations were performed on motion-corrected, dose-weighted micrographs. The thickness measurements in Table 1 are computed with CTFFIND5^31^. Template matching was carried out with an in-house PyTorch-based package, using the same PP7 VLP cryoEM density as the template for both xLPP-on and xLPP-off micrographs.

The 2D Fourier transform of each micrograph was multiplied by the product of (i) a B-factor envelope exp(−B∣k∣^2^/4) with B=100 Å², (ii) a radial band-pass mask (high-pass 800 Å, low-pass 10 Å), (iii) a spectral-whitening filter computed from the rotational power spectrum of the micrograph, and (iv) the contrast transfer function (CTF). CTF parameters used the defocus values estimated by cryoSPARC, with an additional 73° phase shift applied for xLPP-on data.

Template projections were generated on an SO(3) grid at 8° angular resolution. Each preprocessed micrograph was cross-correlated against every rotated projection, yielding at each pixel the best-matching rotation index and its corresponding raw score. Particle picks were obtained by peak detection above a manually optimized threshold, tuned per image to maximize true detections (threshold 9.5 for xLPP-off, 20 for xLPP-on).

For each pick, the pose was further refined by a local search over SO(3) neighbours and 2D real-space translations. For each pose candidate, the shell-wise Fourier ring correlation (FRC) was computed between the template Fourier transform and the CTF-applied image Fourier transform as:

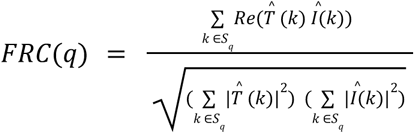

where *S*_*q*_ is the q-th radial Fourier shell, *T* (*k*) is the template projection Fourier transform at the refined rotation, *I*(*k*) is the CTF-applied micrograph particle crop Fourier transform at the refined translational shift. For computing the FRC, only CTF is applied to the micrograph, without other band-pass filters or B factor envelopes. In Fig.5d, mean of the FRCs for (n=41, xLPP-on; n=46, xLPP-off) was plotted as the central curve and the dispersion was shown as a shaded band of the standard error of the mean (±SEM = σ/*N*, where σ is the standard deviation).

### Tilt series collection and tomogram reconstruction

xLPP-on tilt series were acquired with Leginon^25,26^, at 0.465 Å/px in superresolution mode as bidirectional tilt series from 0° to –27° and then 0° to 27° with 3° tilt increments and 4.47 e/Å^2^ per tilt. For every tilt, auto electron-laser alignment is performed at an off-site tracking location. The xLPP was operating at powers corresponding to 90° phase shift, and CTF fit measured 1.5 µm defocus and 72.6° phase shift at 0° tilt. Defocus is 1.39 ± 0.08 µm (mean ± standard deviation) and phase shift is 74° ± 6° for 11 out of 19 total tilts where CTF fitting confidence is better than 10 Å.

Frames were acquired as tifs with five frames from the same microscope and camera as the xLPP-on single particle datasets. Aretomo^32^ pre-processing was performed for estimation of CTF and phase shift from the xLPP, with motion-correction omitted. IMOD^33^ patch-tracking was used for tilt series alignment and the reconstruction of weighted back-projection tomograms at a pixel size of 7.44 Å/px (bin 8x). No denoising or additional filtering was applied for Extended Figure 18 and Movie 1.

## Data availability

The apoferritin maps are deposited on EMDB, with entries specified in Extended Table 2. The paired *E. coli* images with xLPP-on and xLPP-off will be available on EMPIAR upon publication. The through-focus series with xLPP on and xLPP off on amorphous carbon (Fig. 3 and Extended Data Fig. 5) will be available on Zenodo upon publication. The xLPP apoferritin tomogram is available on the cryoET data portal with dataset ID DS-10496.

## Acknowledgments

The authors thank J. Dickerson for xLPP CTF modeling and enlightening discussions on B factors and 2D template matching. The authors thank D. Serwas, H. Siems, N. S. Hill, M. Ali, for the apoferritin and the *E. coli* samples. The authors appreciate K. Z. Zhao’s help on 2D template matching analysis. The authors thank the Biohub high-performance computing (HPC) team and IT staff S. Kaul, V. Kumar, N. Soderquist, W. Law and K. Greer for their computational and IT support. The authors thank U. Ermel for helping publish the xLPP tomogram on the cryoET data portal. The authors thank M. Mouhab, A. Zhang, A. Leshchenko, and K. Gross from Applied Precision Design for their contribution to the design and assembly of the crossed laser phase plate and the laser delivery and optical monitoring systems.

## Author contributions

Y.Y., P.K.O., M.K., D.B., E.C. J.T.Z., H.M., B.C. and D.A. prepared the manuscript with contributions from all co-authors. P.N.P., J.T.Z., J.A. and H.M. conceived the theoretical design for the crossed laser phase plate (xLPP). P.K.O., A.T., D.R., M.D, P.N.P., J.T.Z., N.P., J.A., H.C., J.R, A.S., L.M., C.S.P., H.M., B.C., and D.A. contributed to the engineering design and implementation of the crossed laser phase plate. P.K.O., A.T., and D.R. aligned the optics for the xLPP cavity. B.B. and H.M. contributed to the design of the new LPP module in the TFS Krios. N.P., J.T.Z., J.A., P.N.P., L.M., P.K.O., A.S., H.M. and D.A. contributed to the xLPP optical cavity control systems. P.K.O, A.C., Y.Y and E.C. designed, collected, and analyzed data for xLPP characterization, including determination of the waist parameters of intracavity beams, cavity stability and drift experiments. E.M. and N.P. contributed to initial characterization of cavity stability. A.C., E.M., Y.Y., P.K.O., E.C., P.N.P., J.T.Z., B.B., W.H., B.C. J.A., J.R., and D.A. contributed to xLPP-related microscope alignment and automated data collection procedures. Y.Y., P.K.O., A.C., B.C. and D.A. designed, collected and analyzed data for characterizing the xLPP phase shift on amorphous carbon. E.M., M.K. and M.P. prepared the apoferritin and the *E. coli* samples. E.M., A.C., Y.Y., M.K. and P.K.O. designed, collected and analyzed data demonstrating sub-2 Å resolution apoferritin with xLPP on and off. Y.Y., A.C., M.P., D.K. and M.K. designed, collected and analyzed data for *E. coli* imaging with xLPP and 2D template matching. Y.Y., D.B, S.Z., A.C. and P.K.O. contributed to CTF estimation and simulation for xLPP data. A.C. designed and performed the xLPP tomography collection, J.H. and Y.Y. processed the tomogram.

## Extended Data

**Extended Data Fig. 1.**
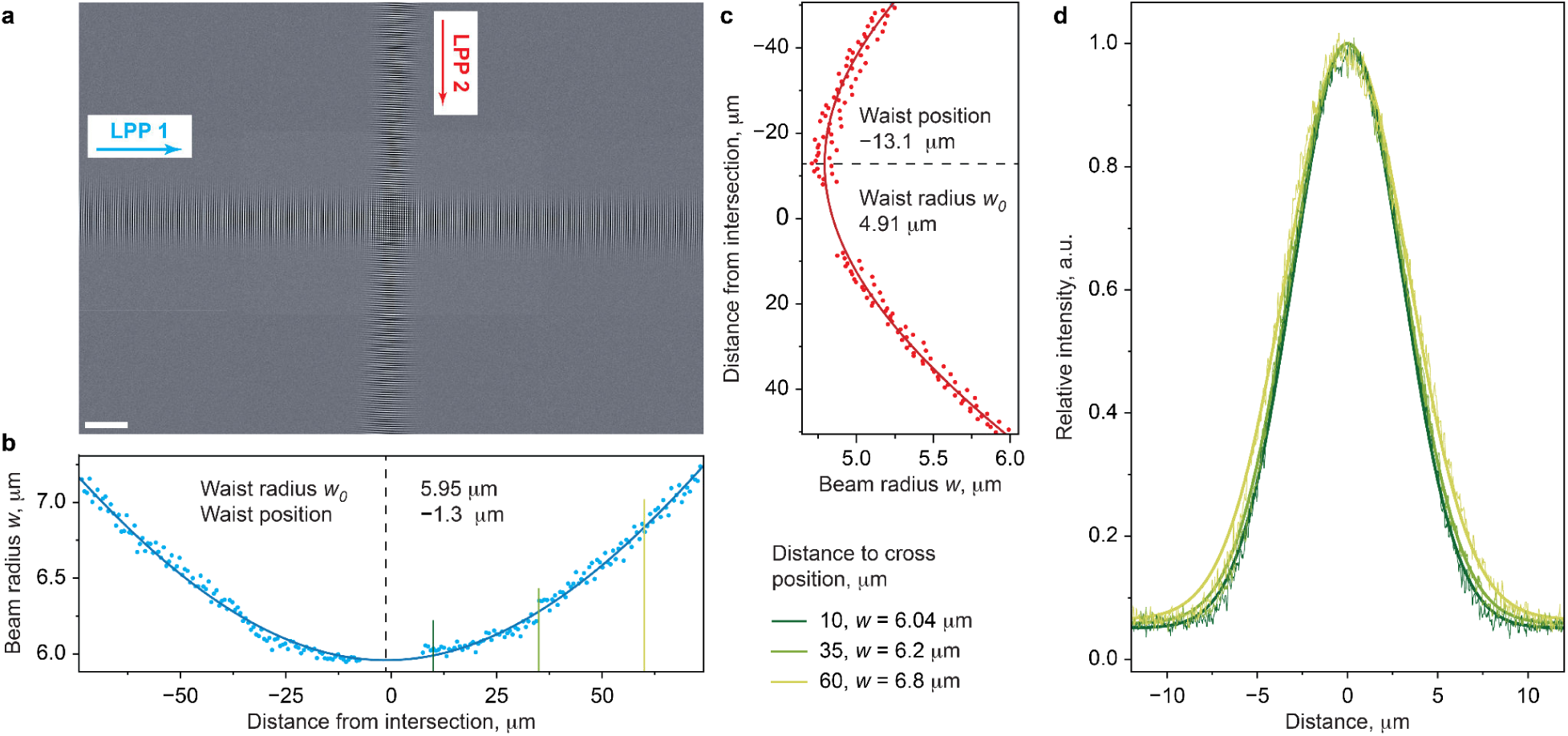
| Characterization of the beam waists for two optical cavities. **a**, Montage image of the Ronchigram of the intersection of two standing waves. **b,c,** Beam radii (1/e^2^) of the standing waves as the function of the distance from the intersection between the beams. The data were fitted with a formula for a Gaussian beam to determine waist positions and waist radii of the beams. The radii were determined from Gaussian fits of the profiles taken across each beam at different distances from the intersection point between the beams. **d,** Representative Gaussian fits (thick lines) of individual profiles (thin lines) taken at different distances to the cross position, as indicated in (**b**).

**Extended Data Fig. 2.**
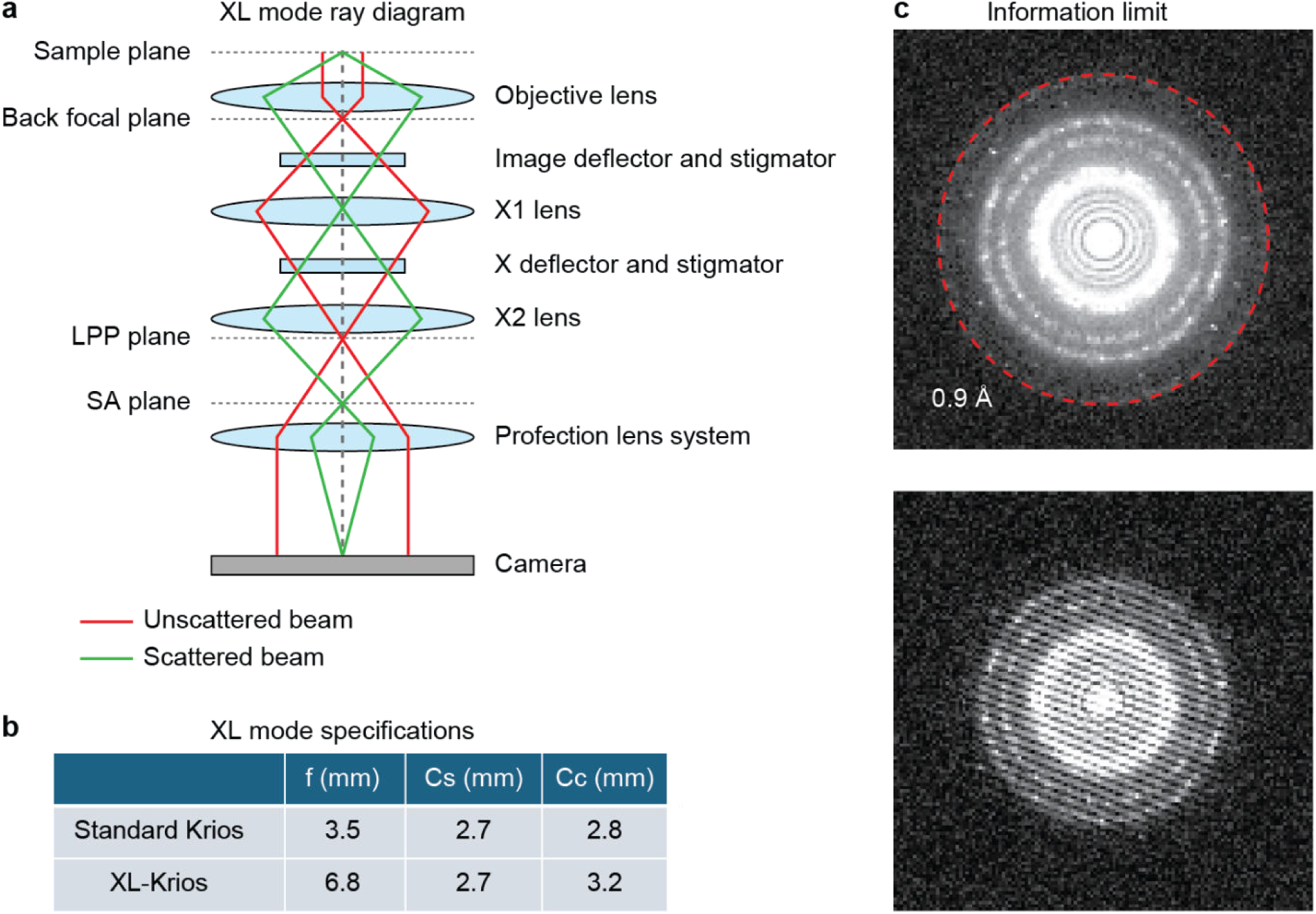
| Optical design and performance of the XL-Krios in HM-XL mode. **a**, Ray diagram of the HM-XL optical configuration, showing the additional transfer lens modules (X1 and X2 lenses) used to project the back focal plane to the phase plate plane. **b,** Comparison of key optical parameters between a standard Krios and the XL-Krios. The modified configuration increases the effective focal length (F) and the chromatic aberration coefficient (Cc) while maintaining the comparable spherical aberration coefficient (Cs). **c,** Information limit assessment using a cross-grating grid. The power spectrum shows signals extending to ∼0.9 Å (top), and corresponding Young’s fringes (bottom), showing information limit at 0.9 Å.

**Extended Data Fig. 3.**
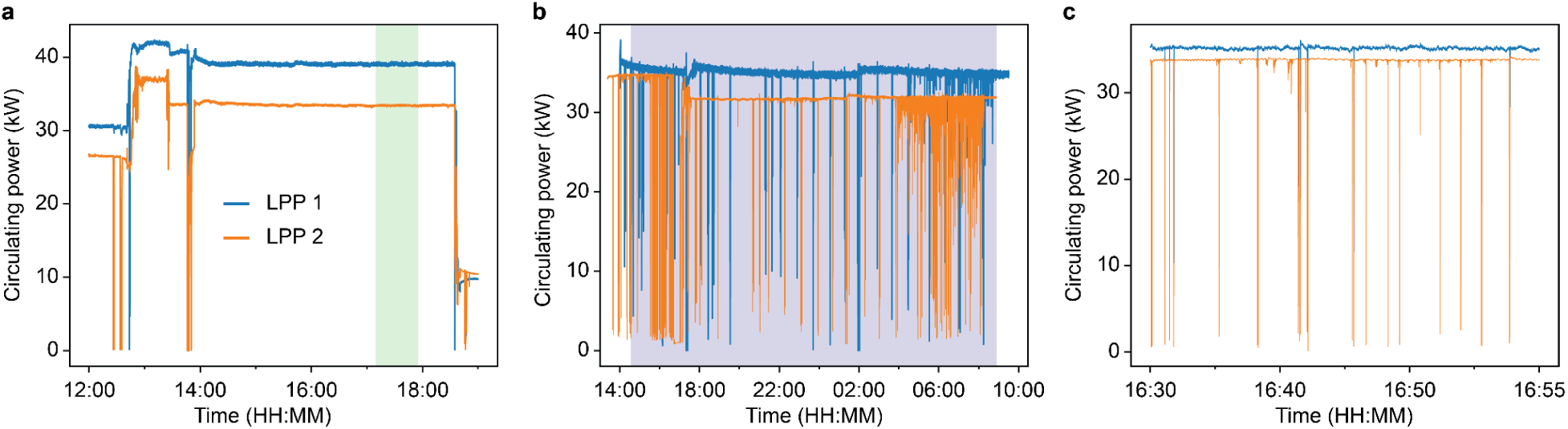
| Cavity power stability during the experimental data acquisition. Circulating powers in both cavities of the xLPP during the experiment presented in Fig. 2a (**a**) and Ext. Data Fig. 11 xLPP-on (Dataset B1) (**b**). The shades demonstrate the actual times of the data acquisition. **c**, Short term stability during the SPA data acquisition.

**Extended Data Fig. 4.**
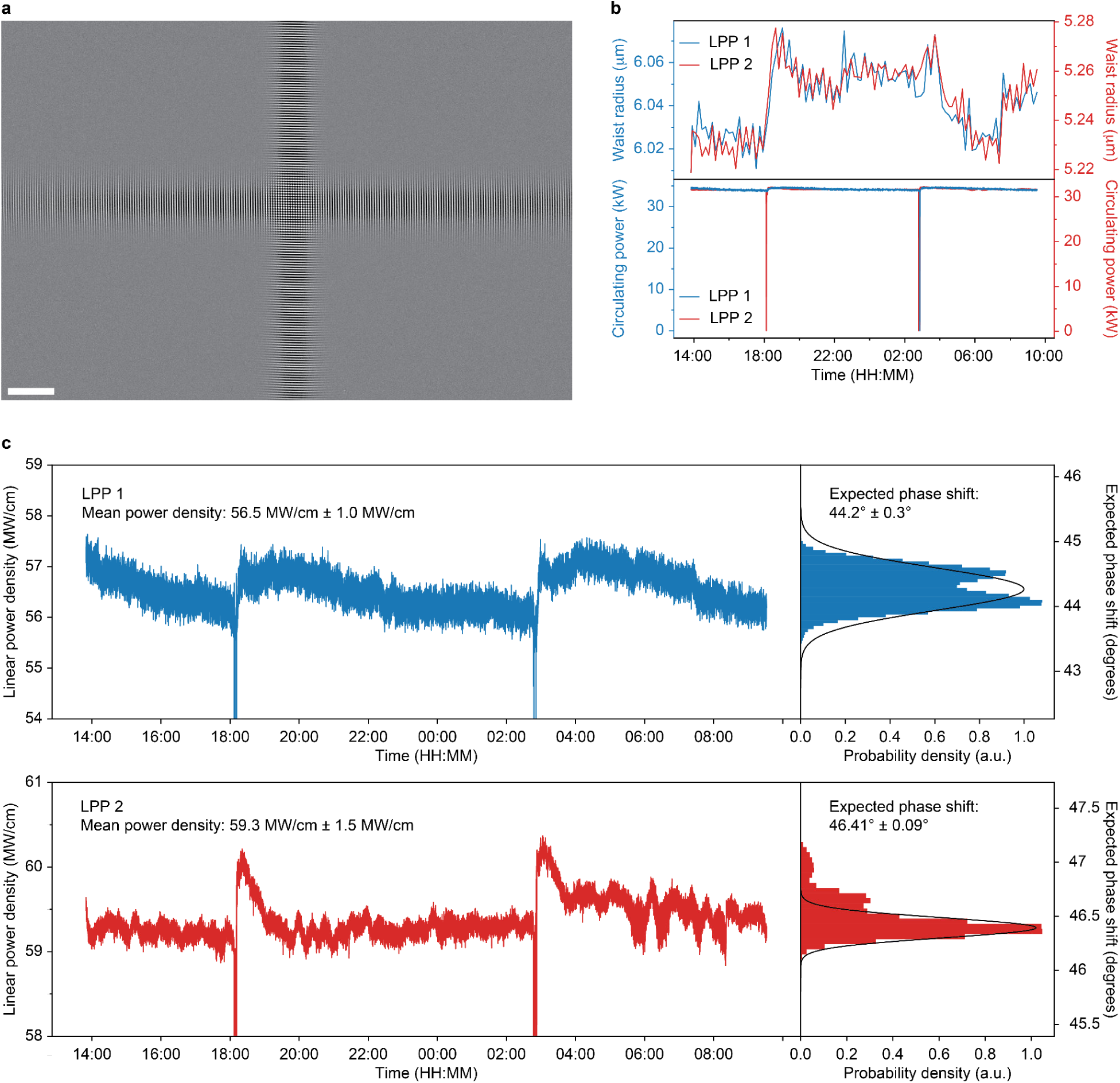
| Characterization of the laser beam stability. **a**, Typical micrograph of the standing waves that was used to determine the beam waists at the intersection point. Scale bar, 10 μm. **b**, Changes in the beam radius at the intersection point and extracted from a series of Ronchigrams (top) and the laser circulating power (bottom). **c**, Intracavity circulating power. **d**, Power density calculated as the ratio of the circulating power from (**c**) and the beam waist radius in (**b**). The histograms demonstrate the estimated phase shift with the Gaussian fit of the data.

**Extended Data Fig. 5.**
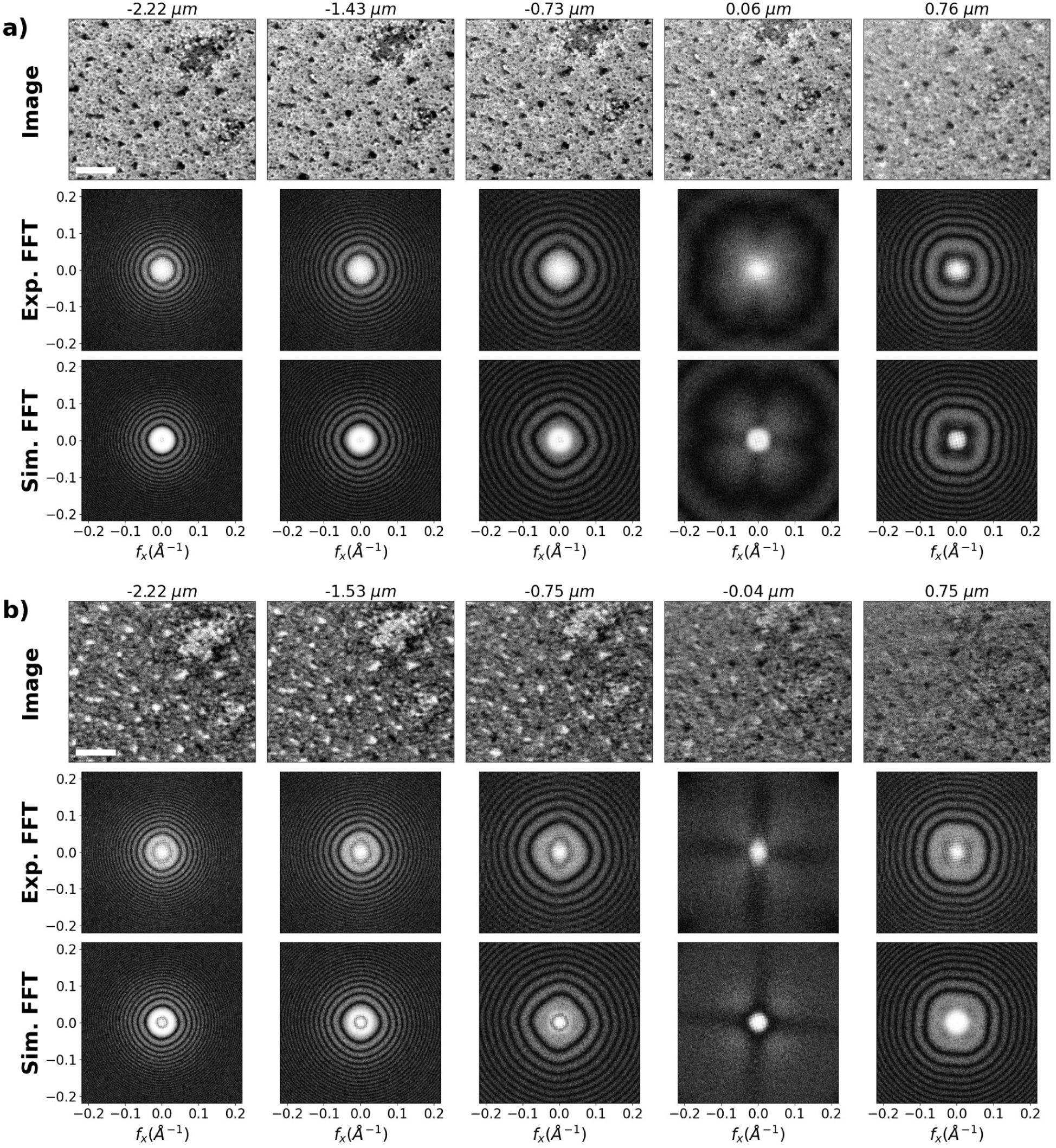
| Experimental and simulated FFTs for xLPP images on amorphous carbon. Representative micrographs (top rows), experimental FFTs (middle rows), and FFTs of simulated images modulated with a xLPP CTF model^24^ (bottom rows). **a,** The top panel corresponds to the electron beam aligned to the antinodes of the xLPP standing waves and **b,** the bottom panel corresponds to the nodes. Scale bars, 50 nm. A range of defocus values was recorded. At the antinodes, the FFTs (log power spectra) show the expected contrast enhancement at low spatial frequencies. At the nodes, scattered electrons at certain spatial frequencies are still phase-shifted by the laser standing wave, producing low-frequency contrast enhancement with inverted contrast. Defocus values were measured for underfocus data and extrapolated for on-focus and overfocus images, as the defocus lens step size is kept constant throughout the series. The discrepancy observed at on-focus and overfocus is likely attributable to lower confidence in CTF estimation under these conditions. The power spectra of the simulated images reproduce the characteristic fourfold pattern arising from the dual-laser configuration, instead of the twofold pattern from previously reported single-laser LPP data.

**Extended Data Fig. 6.**
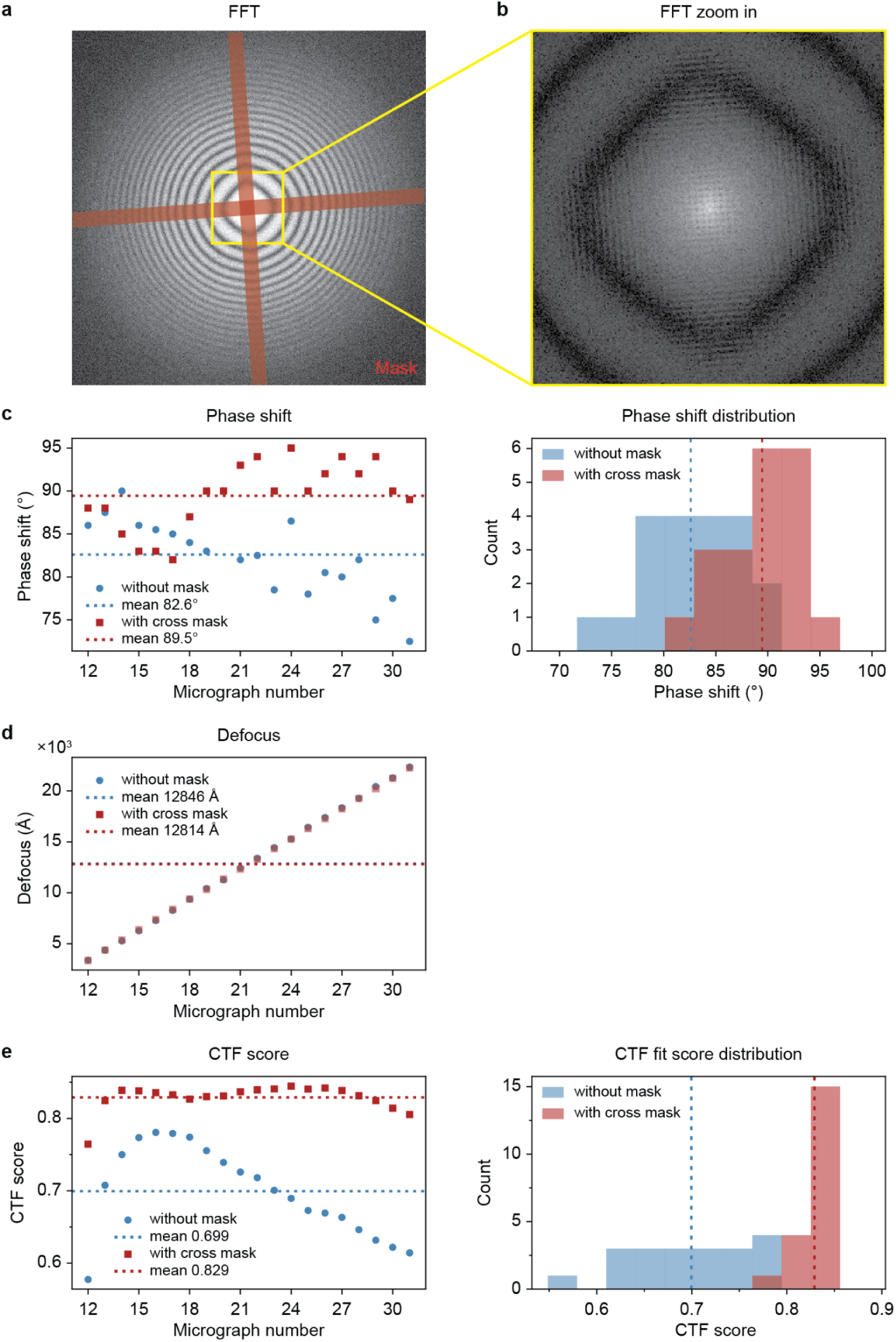
| Laser standing wave impact on CTF estimation. **a**, Representative FFT with the cross-shaped mask (red) overlaid, used to block the laser standing wave signal prior to CTF fitting. **b,** Zoom-in of the central FFT region (yellow box in (**a**)) showing the Thon rings used for fitting. **c,** Phase shift estimates with and without the cross mask across micrographs (left) and their distributions (right); masked fits yield a mean phase shift of 89.5° compared to 82.6° without the mask. **d,** Corresponding defocus estimates with and without the mask (left), showing negligible difference between the two conditions. **e,** CTF fitting confidence score with and without the cross mask across micrographs (left) and their distributions (right). Fitting with the mask yields a higher confidence score.

**Extended Data Fig. 7.**
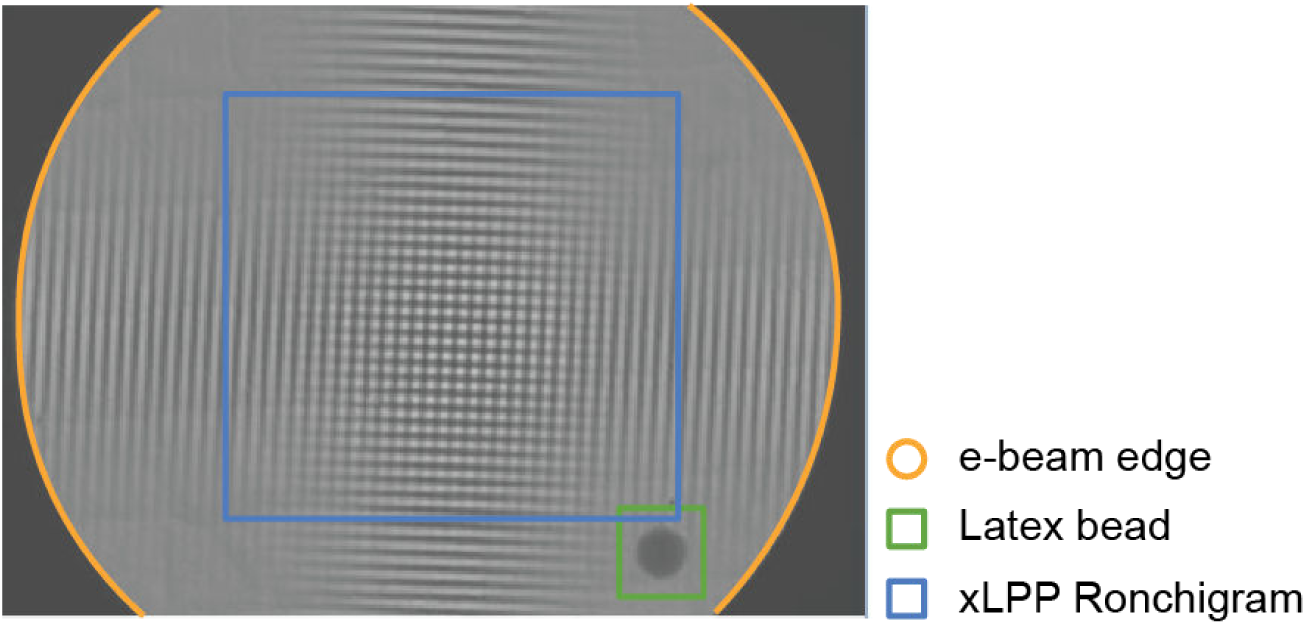
| Example image for temperature induced drift measurements. An example image used for measuring the temperature induced drifts. The orange circle indicates the beam edge, the blue box is the xLPP Ronchigram (far-field interference pattern) and the green box is the latex bead.

**Extended Data Fig. 8.**
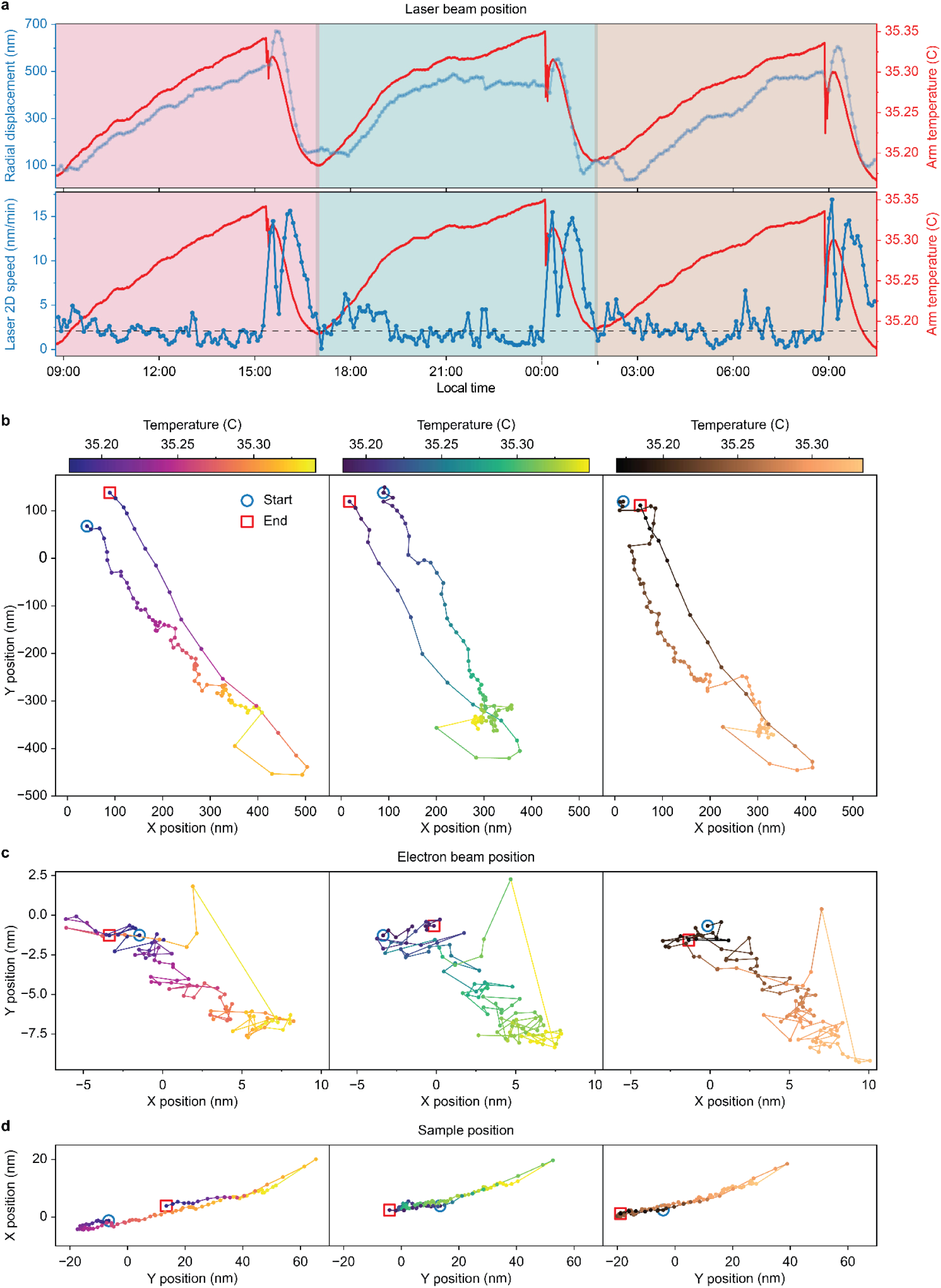
| Temperature induced drifts. **a**, Radial displacement (top, left axis) and corresponding drift (bottom, left axis) of the laser intersection point; support arm temperature is shown on the right axis. The dashed line in the bottom plot represents an average drift ∼7 hours before filling the microscope with liquid nitrogen. **b, c, d,** position of the laser intersection point (**b**), electron beam position (**c**), and a sample position (**d**) at different temperatures of the arm during the time intervals from 08:45 to 17:00, from 16:55 to 01:45, and from 01:40 to 10:30 from left to right.

**Extended Data Fig. 9.**
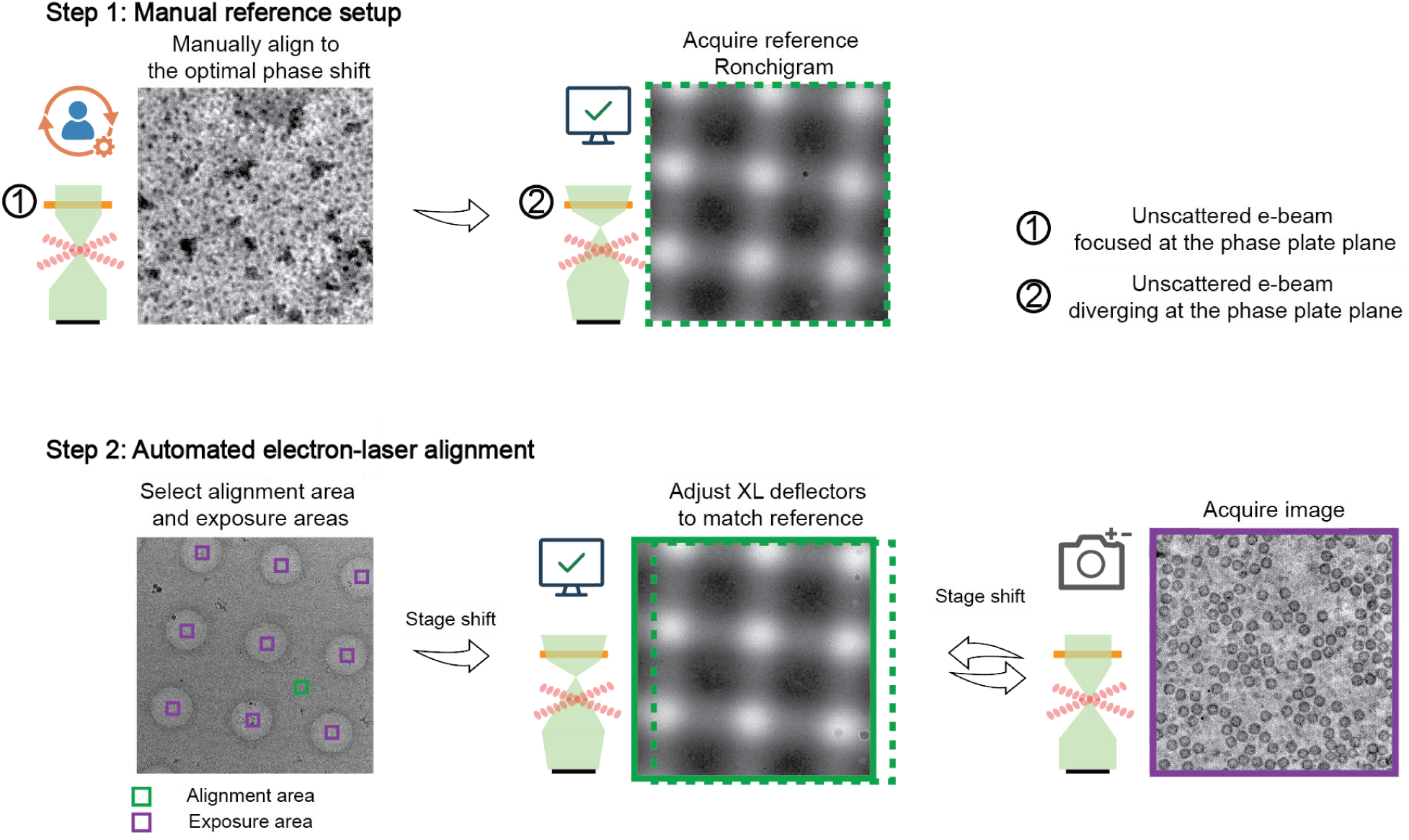
| Schematic of the Leginon-based electron–laser alignment procedure. Alignment (green) and exposure (purple) areas are defined in low-magnification images. The dashed green box marks the reference Ronchigram acquired in Step 1, and the solid green box marks the alignment-check Ronchigram acquired in Step 2. Step 1 (performed once at the beginning of data collection): the user manually adjusts the electron beam to optimize the phase shift under imaging conditions (unscattered beam focused at the phase plate plane); a reference Ronchigram is then acquired at optimal alignment by diverging the beam at the phase plate plane. Step 2 (repeated automatically throughout data collection): the stage is shifted from an exposure area to the alignment area, where an alignment-check Ronchigram is acquired and aligned to the reference Ronchigram by adjusting the XL deflectors; the stage is then shifted to the next exposure area, and the unscattered beam is focused at the phase plate plane for image acquisition. The illustration schematic diagrams are adapted from reference ^34^ figure 10.7.

**Extended Data Fig 10.**
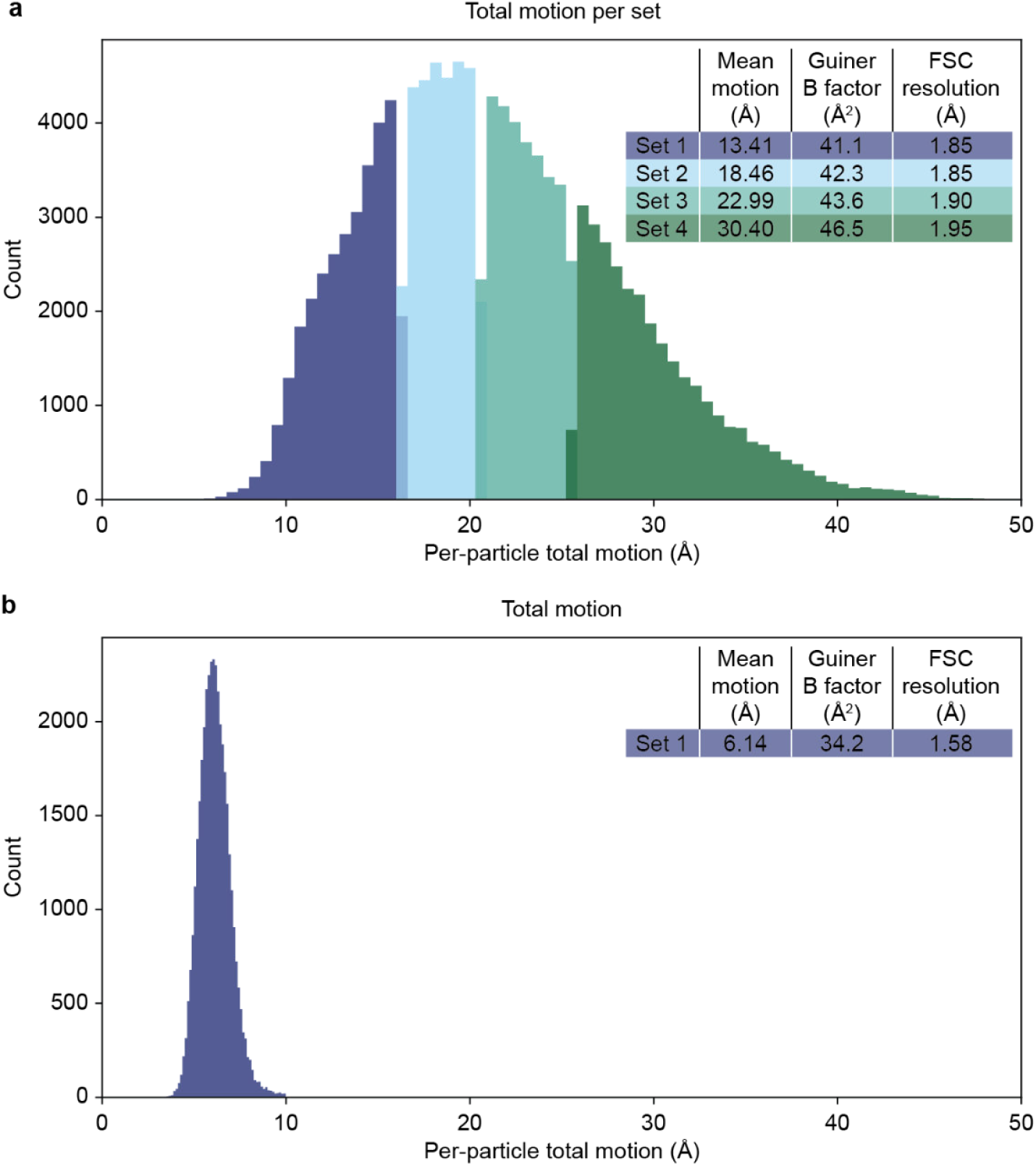
| Resolutions and Guinier B-factors for particle sets with different total per-particle motion. **a**, Particles from dataset A (Fig. 4b,c) were divided into four equal-sized bins by per-particle total motion. The Guinier B-factor and FSC resolution of independent reconstructions from each bin are reported in the inset; both degrade with increasing per-particle total motion. **b,** Reference dataset acquired on a standard (non-LPP) Krios with conventional beam–image shift data collection, on Au R1.2/R1.3 grid type, showing substantially smaller per-particle motion together with a lower B-factor.

**Extended Data Fig. 11.**
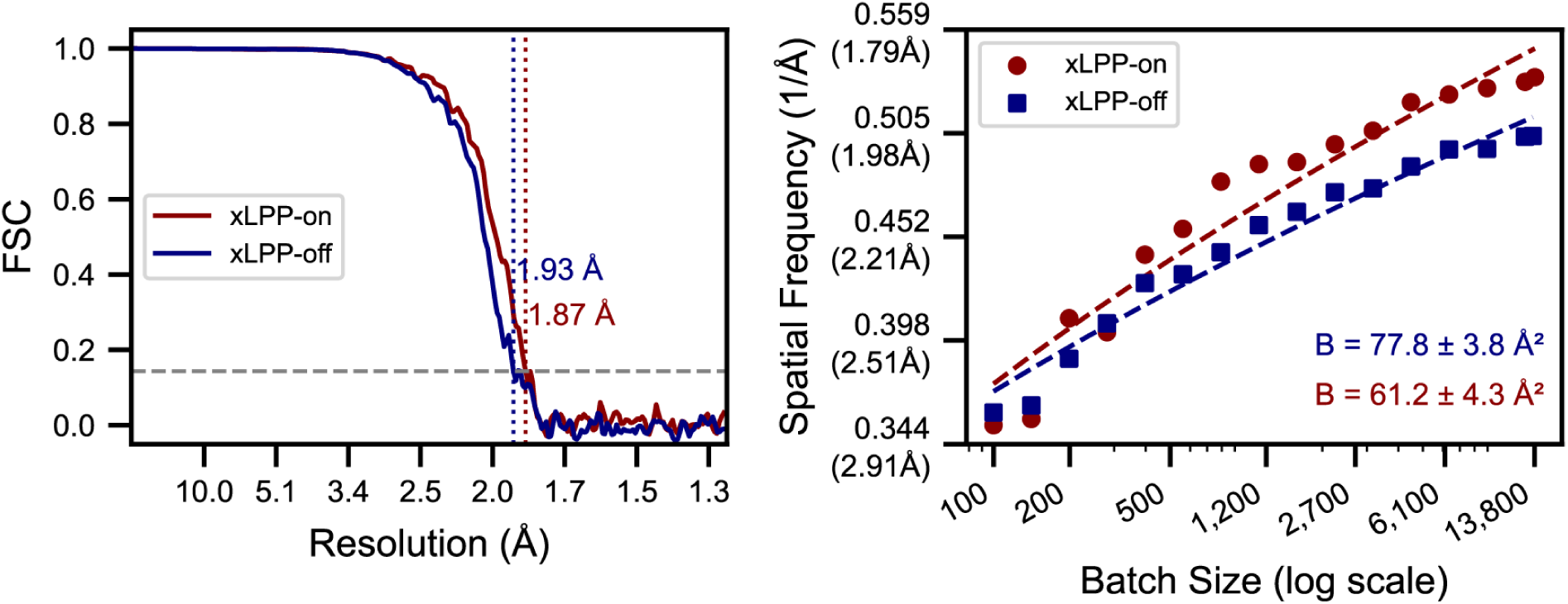
| Apoferritin single particle analysis xLPP on and off. Fourier shell correlation (FSC) for xLPP-on (red) and xLPP-off (blue) reconstructions, yielding resolutions of 1.87 Å and 1.93 Å at the 0.143 criterion, respectively. Rosenthal–Henderson B-factor plots, yielding B-factors of 61 Å² ± 4 Å² (xLPP-on, red) and 78 Å² ± 4 Å² (xLPP-off, blue). Resolutions were determined at the gold-standard FSC cutoff of 0.143.

**Extended Data Fig. 12.**
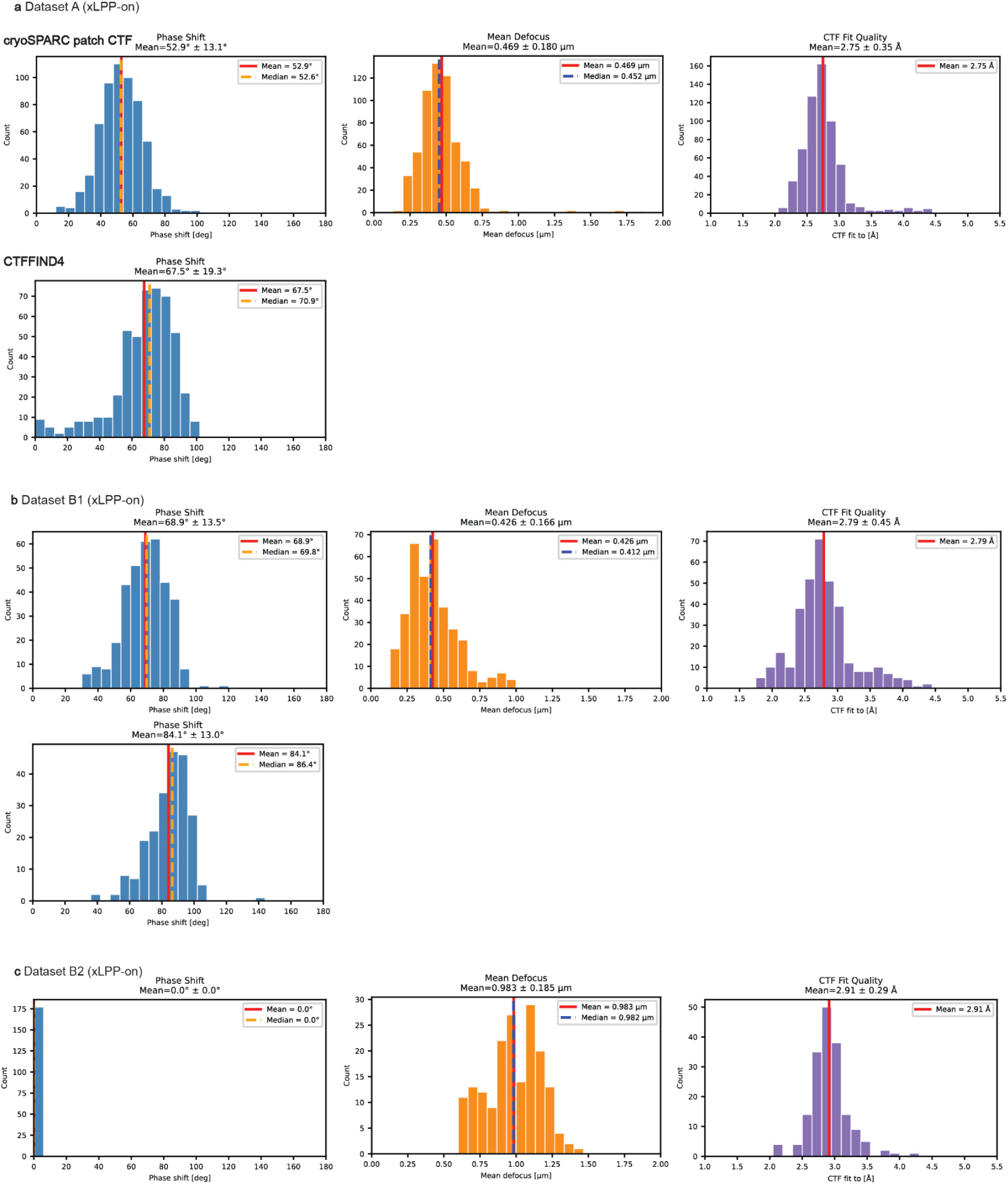
| Statistics of apoferritin SPA data in Fig. 4 (Dataset A) and Ext. Data Fig. 11 (Dataset B1 and B2). The median, mean, and standard deviation for phase shifts, defocus, astigmatism and CTF fit resolution from cryoSPARC. Additionally, for Dataset A and B1, the phase shifts measured by CTFFIND4 are also reported.

**Extended Data Fig. 13.**
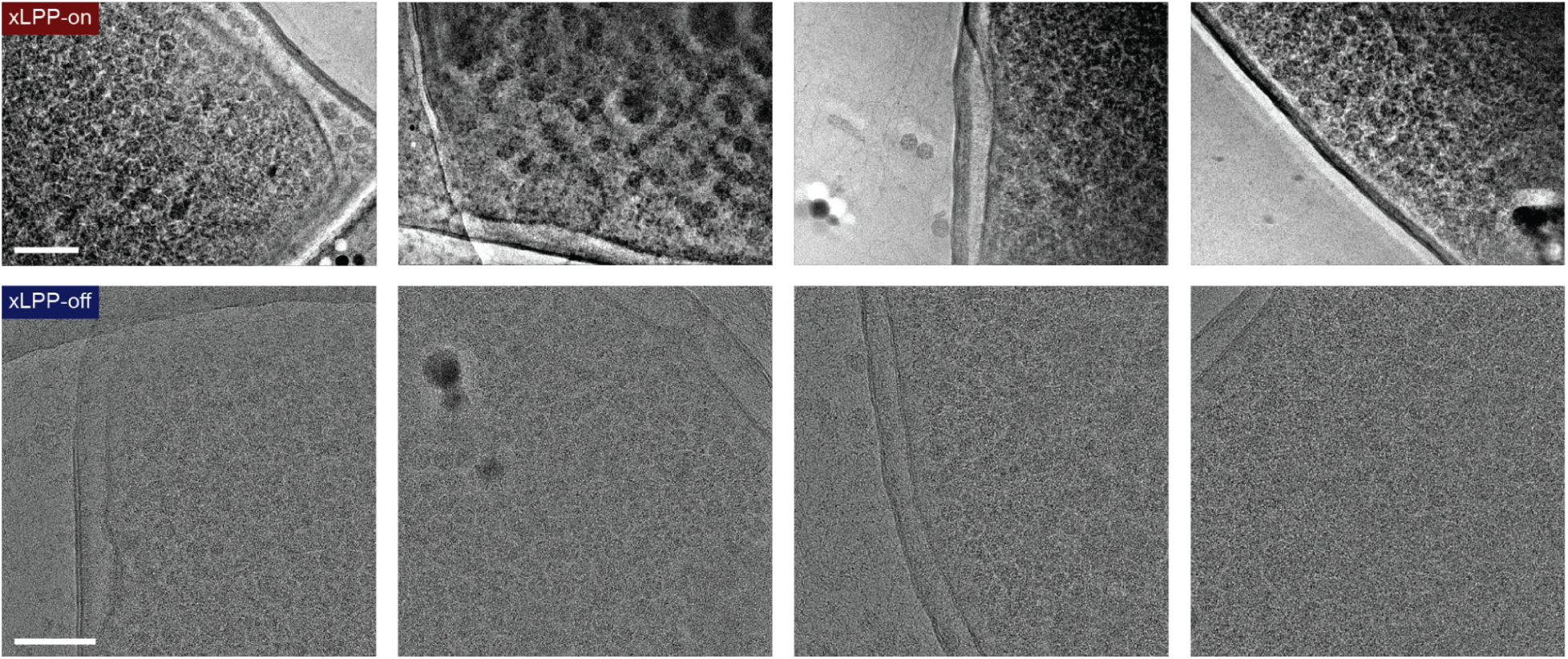
| Gallery of images on *E. coli* cells overexpressing PP7 VLPs with xLPP-on and xLPP-off. The plotted grayscale uses the same threshold percentiles. Top row defocus (from left to right):0.9 µm, 1.0 µm, 0.7µm, 0.7µm; top row phase shifts measured from CTF fitting: 80.8°, 58.5°, 79.3°, 92.4°. Scale bar, 100 nm.

**Extended Data Fig. 14.**
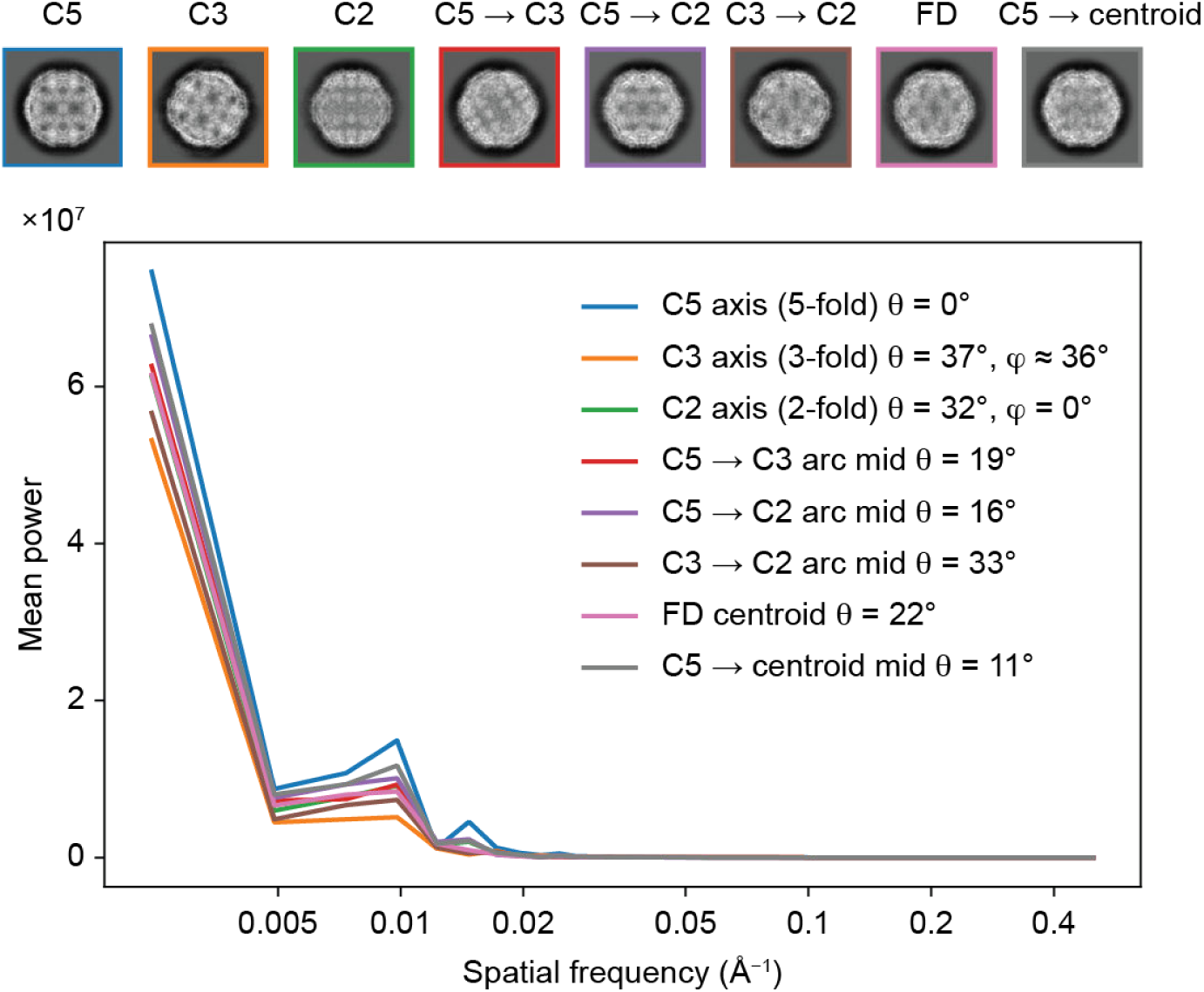
| Signal distribution variation for PP7 VLPs at different projection orientations. Projection images and radial power spectra of the PP7 VLP template across the icosahedral asymmetric unit. Each panel shows the real-space projection (top) at different orientation and the corresponding radial average of the power spectrum (bottom). Projections along different orientations have different signal distributions. This orientation dependence is pronounced for the peak at ∼1/100 Å^-1^, and the range from 1/100 to 1/50 Å^-1^ spatial frequency range, where structural features such as the capsid shell thickness, inter-subunit spacing, and surface spikes contribute strongly to the power spectrum. This orientation-dependent signal is intrinsic to the particle geometry and is independent of CTF.

**Extended Data Fig. 15.**
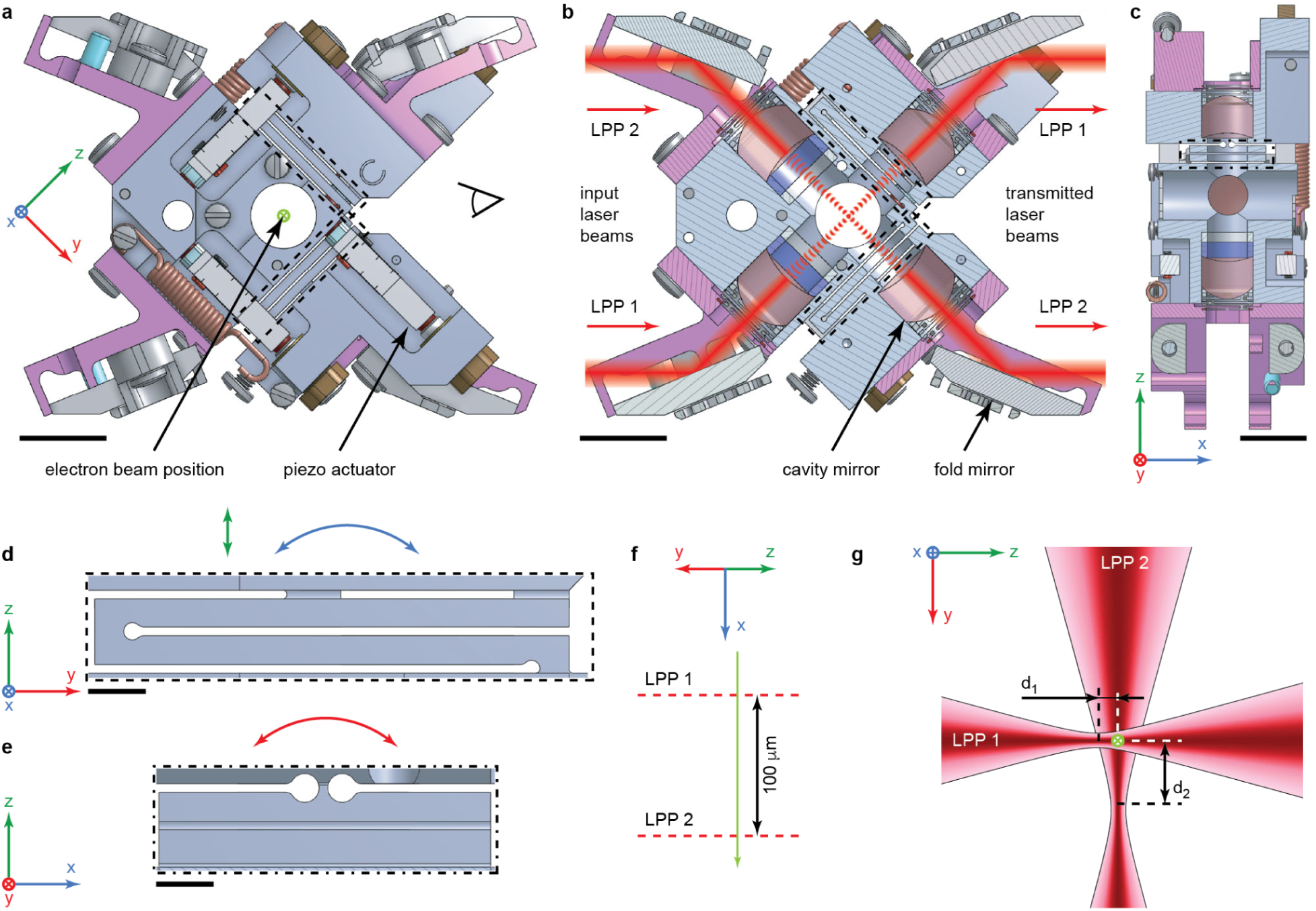
| xLPP flexure design. **a, b**, General view (**a**) of the cavity without the protective cover and a cross section (**b**) of the cavity perpendicular to the x-axis. The electron beam propagates along the x axis. The eye sign indicates the orientation of the schematic in (**f**). Scale bars, 10 mm. **c,** Cross section of the cavity perpendicular to the y axis. Scale bar, 10 mm. Side (**d**) and front (**e**) views of the xLPP flexure marked with dashed rectangles in (**a, b**) and (**c**), respectively. The arrows indicate the directions of flexure motion for the cavity alignment. Scale bars, 2 mm. **f,** Schematic of the relative position of the laser beams in the electron beam propagation direction. **g,** Schematic representation of the central part of the xLPP, d_1_ and d_2_ indicate the distance from the electron beam to the waists of intracavity laser beams.

**Extended Data Fig. 16.**
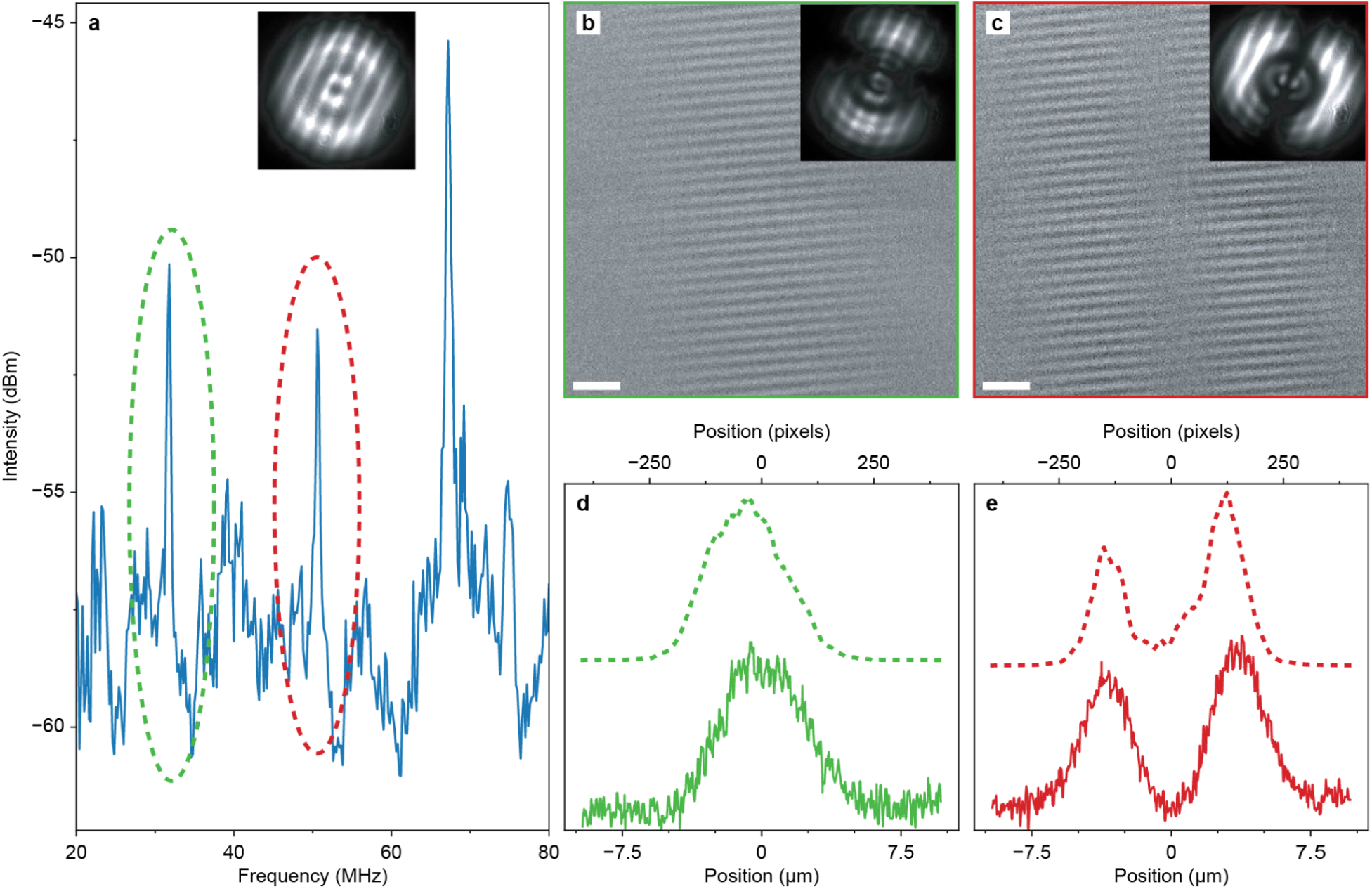
| Effect of the optical cavity astigmatism on the accumulated phase shift in xLPP. **a**, Transverse mode spectrum of the optical cavity LPP2. The inset shows an image of the fundamental mode as recorded with a camera located at a distance of ∼300 mm from the output flange of the microscope. The dashed ovals indicate two peaks around 32 MHz (corresponds to NA of 0.071 and waist radius of 4.75 μm) and 51 MHz (corresponds to NA of 0.056 and waist radius of 6.0 μm) corresponding to TEM_01_ and TEM_10_ cavity modes. **b,c,** Ronchigrams of the laser standing waves locked to the first-order cavity modes. The insets show the images of the corresponding modes recorded at the output of the optical cavity. Scale bars, 2 μm. **d,e,** Profiles extracted from rectangular regions of the Ronchigrams taken in the direction perpendicular to the standing wave propagation (solid lines) and the corresponding camera images (dashed lines) for TEM_01_ and TEM_10_ cavity modes from (**b**) and (**c**), respectively.

**Extended Data Fig. 17.**
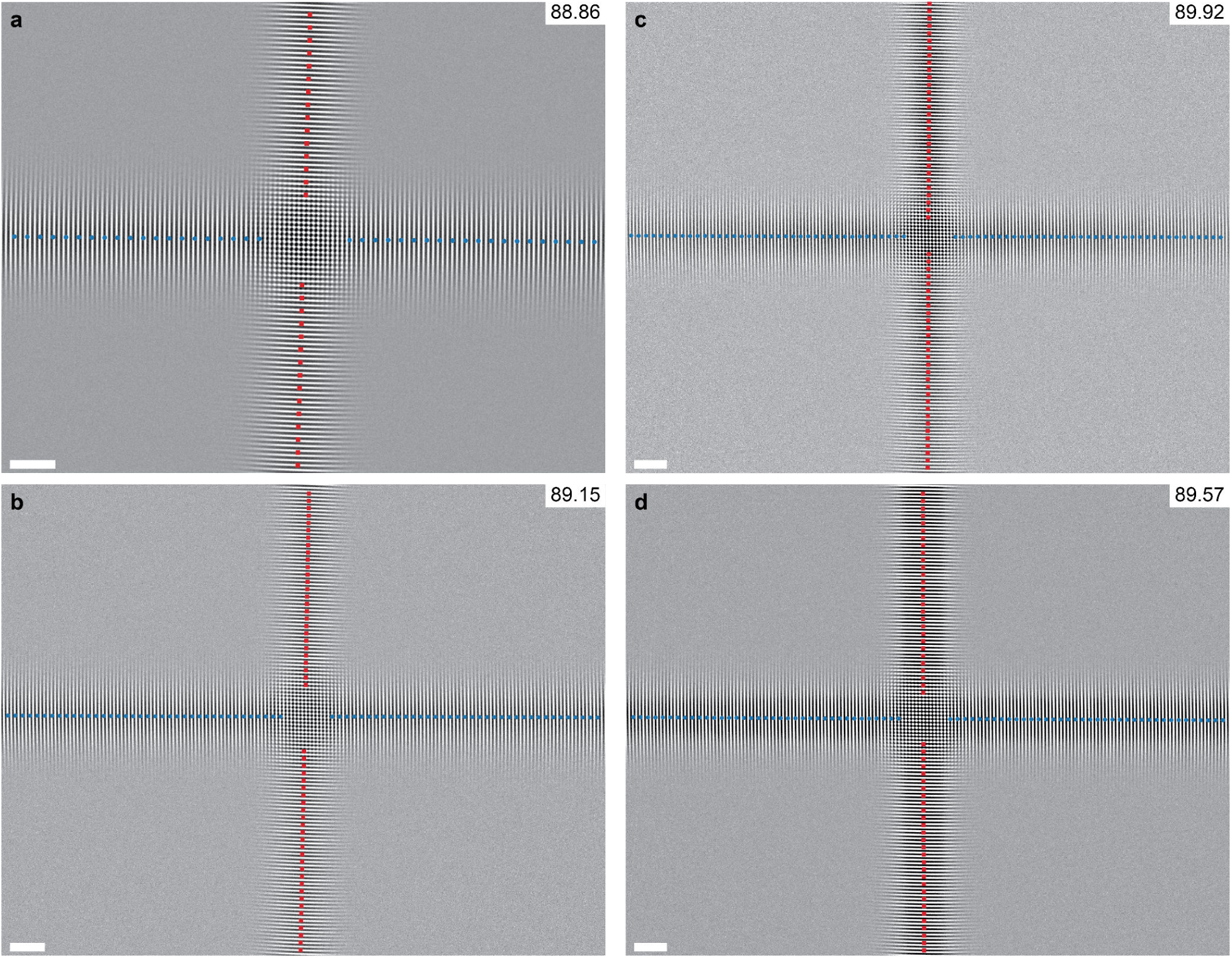
| Determination of angle between optical standing waves. **a-d**, Ronchigrams of the intersection of two standing waves recorded during different experimental sessions. Blue dots and red squares correspond to the centres of Gaussian profiles taken across the standing waves for LPP 1 and LPP 2, respectively. The angles between the lines are shown in the top right corners. Scale bars, 5 μm.

**Extended Data Fig. 18.**
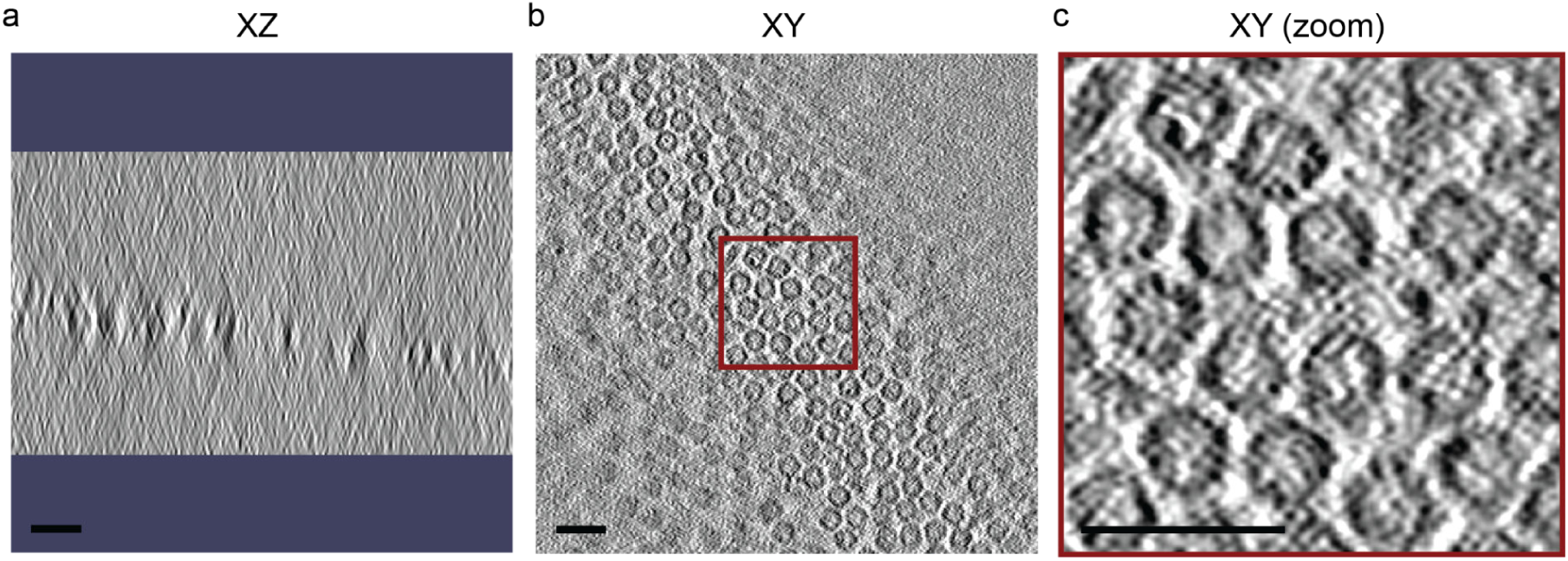
| xLPP-on tomogram of apoferritin. **a-b**, Slices of tomogram in the XZ and XY planes, respectively. The angles between the lines are shown in the top right corners. **c,** enhanced view of central region in **b**. Scale bars, 25 nm. The xLPP was operating at powers corresponding to 90° phase shift, and CTF fit measured 1.5 µm defocus and 72.6° phase shift at 0° tilt. Defocus is 1.39 ± 0.08 µm (mean ± standard deviation) and phase shift is 74° ± 6° for 11 out of 19 total tilts where CTF fitting confidence is better than 10 Å.

**Movie 1 | xLPP-on tomograms of apoferritin.** Slices through the tomogram in the XY plane. Scale bar, 25 nm.

## References

1. Nakane, T. et al. Single-particle cryo-EM at atomic resolution. Nature 587, 152–156 (2020).

2. Tegunov, D., Xue, L., Dienemann, C., Cramer, P. & Mahamid, J. Multi-particle cryo-EM refinement with M visualizes ribosome-antibiotic complex at 3.5 Å in cells. Nat. Methods 18, 186–193 (2021).

3. Glaeser, R. M. How good can cryo-EM become? Nat. Methods 13, 28–32 (2016).

4. Nogales, E. The development of cryo-EM into a mainstream structural biology technique. Nat. Methods 13, 24–27 (2016).

5. Zernike, F. Phase contrast, a new method for the microscopic observation of transparent objects. Physica 9, 686–698 (1942).

6. Boersch, H. Über die Kontraste von Atomen im Elektronenmikroskop. Z. Für Naturforschung A 2, 615–633 (1947).

7. Dwyer, C. & Paganin, D. M. Quantum and classical Fisher information in four-dimensional scanning transmission electron microscopy. Phys. Rev. B 110, 024110 (2024).

8. Danev, R., Tegunov, D. & Baumeister, W. Using the Volta phase plate with defocus for cryo-EM single particle analysis. eLife 6, e23006 (2017).

9. Danev, R., Buijsse, B., Khoshouei, M., Plitzko, J. M. & Baumeister, W. Volta potential phase plate for in-focus phase contrast transmission electron microscopy. Proc. Natl. Acad. Sci. 111, 15635–15640 (2014).

10. Buijsse, B., Trompenaars, P., Altin, V., Danev, R. & Glaeser, R. M. SPECTRAL DQE OF THE VOLTA PHASE PLATE. Ultramicroscopy 218, 113079 (2020).

11. Danev, R. et al. Routine sub-2.5 Å cryo-EM structure determination of GPCRs. Nat. Commun. 12, 4333 (2021).

12. Turoňová, B. et al. Benchmarking tomographic acquisition schemes for high-resolution structural biology. Nat. Commun. 11, 876 (2020).

13. Hutchings, J. et al. Evaluating the Volta phase plate for improved tomogram alignment in cryo-electron tomography. IUCrJ 13, 260–272 (2026).

14. Müller, H. et al. Design of an electron microscope phase plate using a focused continuous-wave laser. New J. Phys. 12, 073011 (2010).

15. Schwartz, O. et al. Laser phase plate for transmission electron microscopy. Nat. Methods 16, 1016–1020 (2019).

16. Turnbaugh, C. et al. High-power near-concentric Fabry–Perot cavity for phase contrast electron microscopy. Rev. Sci. Instrum. 92, 053005 (2021).

17. Axelrod, J. J. et al. Observation of the Relativistic Reversal of the Ponderomotive Potential. Phys. Rev. Lett. 124, 174801 (2020).

18. Remis, J. et al. Cryo-EM phase-plate images reveal unexpected levels of apparent specimen damage. J. Struct. Biol. 216, 108150 (2024).

19. Axelrod, J. J. et al. Overcoming resolution loss due to thermal magnetic field fluctuations from phase plates in transmission electron microscopy. Ultramicroscopy 249, 113730 (2023).

20. Petrov, P. N. Laser phase plate improves structure determination of small proteins by cryo-EM. Science, to be published.

21. Axelrod, J. J. et al. A thermoelastic limit on the focal intensity in Fabry-Pérot cavities. Preprint at 10.48550/arXiv.2604.07676 (2026).

22. Petrov, P. N., Zhang, J. T., Axelrod, J. J., Olshin, P. K. & Müller, H. Crossed laser phase plates for transmission electron microscopy. Nat. Commun (2026).

23. Zheng, S. GCtfFind. (2025).

24. EuanPyle et al. teamtomo: modular Python packages for cryo-EM and cryo-ET.

25. Potter, C. S. et al. Leginon: a system for fully automated acquisition of 1000 electron micrographs a day. Ultramicroscopy 77, 153–161 (1999).

26. Cheng, A. et al. Leginon: New features and applications. Protein Sci. Publ. Protein Soc. 30, 136–150 (2021).

27. Maki-Yonekura, S., Kawakami, K., Takaba, K., Hamaguchi, T. & Yonekura, K. Measurement of charges and chemical bonding in a cryo-EM structure. Commun. Chem. 6, 98 (2023).

28. Danev, R., Yanagisawa, H. & Kikkawa, M. Cryo-EM performance testing of hardware and data acquisition strategies. Microscopy 70, 487–497 (2021).

29. Cheng, A. et al. High resolution single particle cryo-electron microscopy using beam-image shift. J. Struct. Biol. 204, 270–275 (2018).

30. Schaffer, M. et al. Optimized cryo-focused ion beam sample preparation aimed at in situ structural studies of membrane proteins. J. Struct. Biol. 197, 73–82 (2017).

31. Elferich, J., Kong, L., Zottig, X. & Grigorieff, N. CTFFIND5 provides improved insight into quality, tilt and thickness of TEM samples. eLife 13, (2024).

32. Peck, A. et al. AreTomoLive: automated reconstruction of comprehensively corrected and denoised cryo-electron tomograms in real time and at high throughput. Nat. Methods 1–5 (2026) doi:10.1038/s41592-026-03093-y.

33. Mastronarde, D. N. & Held, S. R. Automated tilt series alignment and tomographic reconstruction in IMOD. J. Struct. Biol. 197, 102–113 (2017).

34. Axelrod, J. J. A Laser Phase Plate for Transmission Electron Microscopy. Preprint at 10.48550/arXiv.2403.10670 (2024).

